# Molecular requirements for transition from lateral to end-on microtubule binding and dynamic coupling

**DOI:** 10.1101/481713

**Authors:** Manas Chakraborty, Ekaterina Tarasovetc, Anatoly V. Zaytsev, Maxim Godzi, Ana C. Figueiredo, Fazly I. Ataullakhanov, Ekaterina L. Grishchuk

## Abstract

Accurate chromosome segregation relies on microtubule end conversion, the ill-understood ability of kinetochores to transit from lateral microtubule attachment to durable association with dynamic microtubule plus-ends. The molecular requirements for this conversion and the underlying biophysical mechanisms are ill-understood. We reconstituted end conversion *in vitro* using two kinetochore components: the plus end–directed kinesin CENP-E and microtubule-binding Ndc80 complex, combined on the surface of a microbead. The primary role of CENP-E is to ensure close proximity between Ndc80 complexes and the microtubule plus-end, whereas Ndc80 complexes provide lasting microtubule association by diffusing on the microtubule wall near its tip. Together, these proteins mediate robust plus-end coupling during several rounds of microtubule dynamics, in the absence of any specialized tip-binding or regulatory proteins. Using a Brownian dynamics model, we show that end conversion is an emergent property of multimolecular ensembles of microtubule wall-binding proteins with finely tuned force-dependent motility characteristics.

## INTRODUCTION

Accurate chromosome segregation involves kinetochore attachment to dynamic microtubule (MT) plus-ends, which drive chromosome oscillations in metaphase and pull sister chromatids apart in anaphase. Although some kinetochores acquire such end-on attachments via a direct capture of the growing MT tips, most initially interact with the MT walls^1^. The resultant lateral configuration is subsequently converted to MT end-binding either by depolymerization of a distal MT segment or by the transport activity of CENP-E, a kinetochore-associated plus-end–directed kinesin^2–6^. After the MT plus-end comes in contact with the kinetochore, the chromosomes motions become coupled to MT dynamics, as tubulins are added or removed from the kinetochore-embedded MT ends^7–9^. The biophysical mechanisms underlying conversion of lateral attachment into dynamic MT end-coupling are not well understood^10^. For example, it is not known whether such conversion requires proteins with distinct MT-binding activities, i.e., those that interact with the MT wall during lateral CENP-E–dependent transport and those that subsequently bind to the MT tip. The key motility characteristics of these molecular components have not been determined, and it previously remained unclear whether their interactions with the MT must be regulated in order to enable their distinct interactions with MT walls vs. ends.

To investigate these outstanding questions, we used reductionist approaches with stabilized and dynamic MTs *in vitro*. Specifically, we sought to determine whether the MT wall-to-end transition via the CENP-E– dependent pathway could be recapitulated by combining CENP-E with various kinetochore microtubule-associated proteins (MAPs). Previous work in cells identified the Ndc80 complex as the major kinetochore MAP responsible for end-on MT coupling^11^. However, past *in vitro* reconstitutions provided little insight into why this protein plays such a central role given that it has no intrinsic MT end-binding activity. Indeed, single Ndc80 molecules bind to MTs *in vitro* and undergo transient diffusion along polymerized tubulins in the straight MT wall, but Ndc80 exhibits no strong preference for polymerizing or depolymerizing MT tips^12,13,14,15^. Consistent with this, Ndc80 has significantly lower affinity for curved tubulin protofilaments than intact MTs^12,16^. Although antibody-induced Ndc80 clusters^13^ and Ndc80-coated microbeads^17^ are capable of tracking the dynamic MT ends, this behavior is not unique among MT-binding proteins, including those that have no role in kinetochore-MT coupling^18^. Thus, it remained unclear whether Ndc80 in combination with kinesin CENP-E is capable of supporting MT end conversion. Here, we show that CENP-E and Ndc80 have finely tuned molecular characteristics enabling them to robustly convert lateral MT attachment into end-coupling in the absence of other kinetochore proteins and regulatory events.

## RESULTS

### Ndc80 exerts molecular friction to the CENP-E motor-driven motility on the MT wall

Full-length CENP-E can walk to MT plus-ends and briefly (< 20 s) maintain MT end-association thanks to a MT-binding domain in its tail region^19^. To determine how Ndc80 effects CENP-E motor motility without the interference of this domain, we used a truncated version of CENP-E that falls off MT tips, but on MT walls it walks and responds to force similarly to the full-length protein^20^. Glass microbeads coated with these motor domains (hereafter referred to as “CENP-E motor”) were also coated with purified Ndc80 “Broccoli” complexes containing the wild-type MT-binding domains (Fig. S1a). Using a laser trap, we captured and brought such a bead to the wall of a taxol-stabilized MT lying on a coverslip (Fig. 1a). The bead’s motility depended strongly on the ratio of Ndc80/CENP-E coating (Fig. 1b). With more Ndc80 present, beads tended to pause at the MT tips, detaching less frequently. However, the beads walked more slowly, and many could not reach the MT plus-ends.

**Figure 1.**
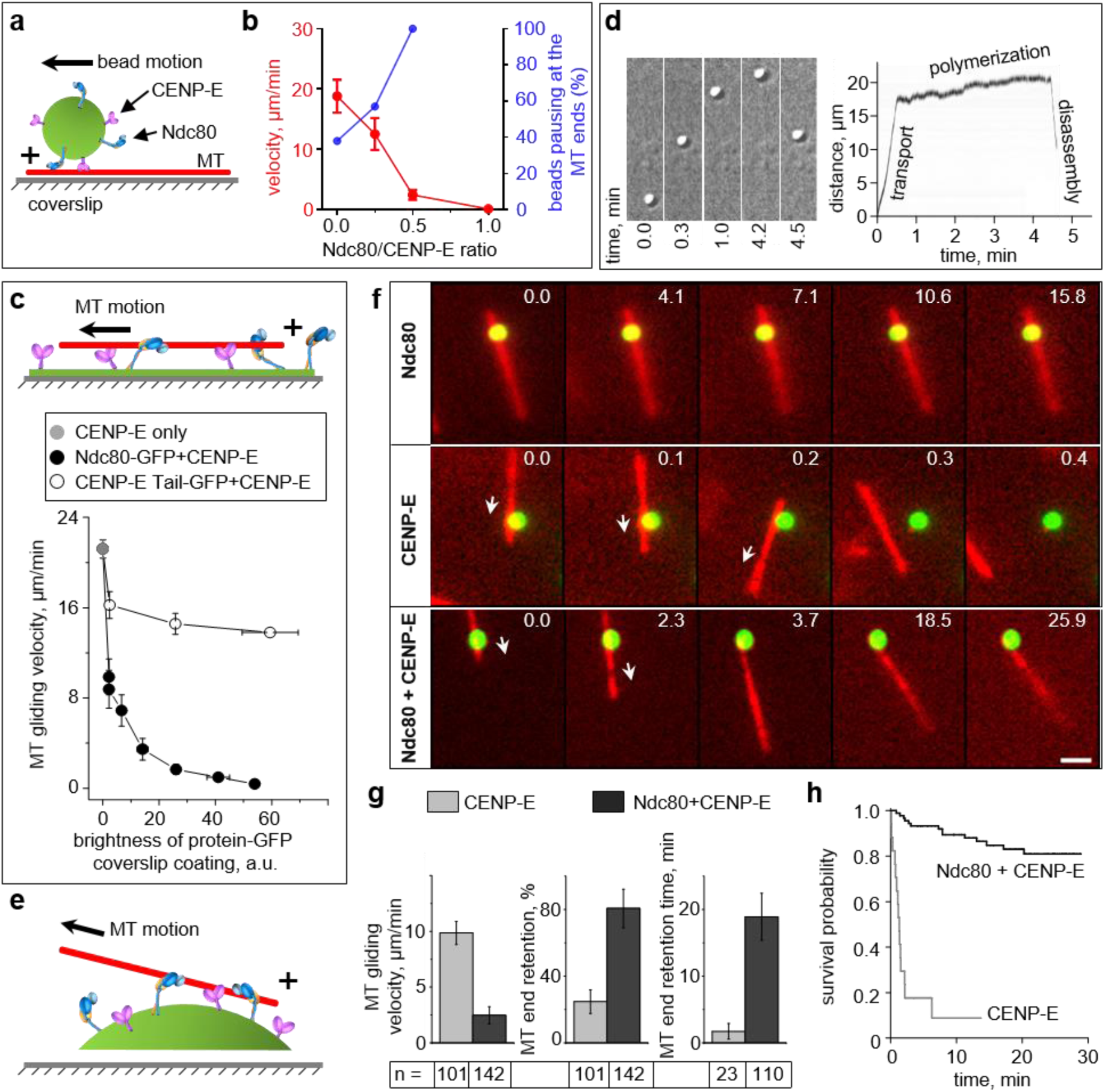
MT wall and tip interactions with a molecular lawn containing a mixture of randomly distributed CENP-E motors and Ndc80 complexes. (a) Experiment with taxol-stabilized MT immobilized on a coverslip and a bead coated with Ndc80 and CENP-E motor. (b) Velocity of bead transport along the MT walls (means ± SEM; red curve plotted using left axis) and attachment at the MT end (blue, right axis) versus the ratio of Ndc80 and CENP-E concentrations used for bead coating. Beads were scored as pausing at the MT tip if they remained attached for at least 2 sec. (c) MT gliding on a coverslip coated with protein mixtures. Top: schematic of coverslip conjugation using tag-binding antibodies, ensuring that MT-binding domains are not sterically inhibited. Bottom: Gliding velocity on coverslips with CENP-E motor and either Ndc80-GFP or CENP-E Tail-GFP versus the protein coating density, as determined by GFP fluorescence. Points are means ± SEM for average velocities from *N*=3 independent trials, each examining n ≥ 54 MTs. (d) Experimental example of a bead walking to the end of the dynamic MT at the velocity of CENP-E motor, continuing in the same direction at the velocity of MT polymerization, and then moving backward when the MT disassembles (MT tip-tracking). Left: time-lapse images acquired with differential interference contrast. Right: kymograph of the entire trajectory. (e) Schematics of the MT wall-to-end transition assay, which uses GMPCPP-stabilized MT seeds and GFP-tagged proteins conjugated to coverslip-immobilized beads. (f) Selected images showing motions of MTs (red) on immobilized beads (green) coated with the indicated proteins. Numbers are time (min) from the start of observation. Arrows show direction of MT gliding. Bar, 3 µm. (g) Quantifications for the wall-to-end transition assay from *N*=4 independent trials for CENP-E motor, and *N*=7 trials for Ndc80+CENP-E motor. Data are means ± SEM for averaged results from each trial. Total number *n* of observed events is indicated under each bar. (h) Kaplan–Meier survival plot for MT end-retention time based on *N* independent trials examining end-attachment for *n* MTs: CENP-E motor: *N*=4, *n*=23; Ndc80+CENP-E, *N*=7, *n*=110.

To investigate the origin of the observed decrease in velocity, we turned to the established “gliding assay,” in which lateral MT transport is driven by a CENP-E motor sparsely attached to the surface of a coverslip (Fig. 1c, S1b). MTs glided at ∼20 µm/min on coverslips coated with CENP-E motor in the Ndc80 absence^21^, but velocity decreased when Ndc80 was conjugated to the same coverslips. Importantly, the Ndc80 and CENP-E motor domains have not been reported to physically interact^22^. Moreover, these proteins were immobilized using different antibodies, and soluble proteins were washed away. Therefore, slow velocity was caused via their mechanical coupling to MTs, rather than direct Ndc80-CENP-E binding. The molecular friction generated by binding of Ndc80 to the MT as it glides under the power strokes of CENP-E slows down the motor. This force-dependent effect is specific to Ndc80, as the MT-binding CENP-E Tail domain caused only a minor hindrance (CENP-E Tail, Fig. 1c), consistent with the tail’s inability to slow down motor domains within the full-length molecule^19^.

Importantly, within the optimal range of Ndc80 and CENP-E coatings on the microbeads, the motor was able to overcome this friction and deliver beads to MT plus-ends. On dynamic MTs growing from coverslip-immobilized MT seeds, upon arrival at the tips the beads slowed down even further and continued to move at the normal rate of MT elongation (Fig. 1d, Fig. S2a-c). When MTs disassembled, the beads moved backward; these beads detached more frequently than during polymerization, but overall their MT end-coupling was maintained for several minutes.

### A pair of only two proteins, CENP-E motor and Ndc80, can enable the transition from wall to end of a stabilized MT

The results obtained using laser-handled microbeads suggested that a combination of CENP-E motors and Ndc80 molecules can support MT end conversion. However, this conclusion was tempered by the observation that MAP-coated beads could roll on the MT surface, a highly non-physiological behavior that would disrupt normal MT end-coupling^19^. Furthermore, the thermal motions of the beads at the ends of MT extensions in this assay complicate thorough analysis and visualization of MT–bead coupling. To overcome these limitations, we modified the assay’s geometry to use coverslip-immobilized microbeads and freely floating fluorescent MTs (Fig. 1e). As expected, beads coated with Ndc80 bound to the walls, rather than the ends, of GMPCPP-stabilized MT seeds (Fig. 1f). MT walls also bound to CENP-E–coated beads in the presence of AMP-PNP, a non-hydrolysable ATP analogue. After ATP was added, the attached MTs glided on the beads (Video 1). Because CENP-E is a plus-end–directed motor, a MT glides with its minus end forward, and the MT plus-end is the last contact point between the MT and the bead. Most MTs detached immediately after their plus-ends arrived at the CENP-E–coated bead, while 24% of ends were transiently retained (1.1±0.3 min; Fig. 1g, S1c), confirming the inability of CENP-E motor domains alone to form lasting attachments to MT tips^19^.

We then supplemented these surfaces with Ndc80 coating that reduced CENP-E gliding velocity 4-fold while allowing the majority of MT plus-ends to arrive at the bead. To estimate the number of Ndc80 molecules that interacted with the MT wall under these conditions, we took advantage of the ability of bead-conjugated Ndc80 to support diffusion of the laterally bound MT (in the absence of CENP-E). We then compared this experimental diffusion rate with the theoretically predicted rate for different numbers of Ndc80 molecules, based on the diffusion rate measured for a single Ndc80 molecule (Supplementary Note, Fig. S3a-c). We estimate that 10–15 bead-bound Ndc80 molecules engaged in MT binding in our end-conversion assay, similar to the number of Ndc80 molecules interacting with one kinetochore MT^23,24^.

Under these conditions, almost 80% of MT plus-ends that arrived at beads containing both proteins remained attached for 18.1±1.2 min (Video 2). The actual end-retention time was even longer, as many end-attachments outlasted a typical experiment. Kaplan–Meier survival analysis, which takes into account the MTs that were still bead-bound at the end of observation, indicated that 80% of MT-end attachments survived for >28 min. On the timescale of mitosis, this constitutes a durable attachment that greatly exceeds results obtained for CENP-E motor alone, with which >80% of MTs detached in <2.5 min (Fig. 1h). Because these beads are immobilized, rolling could be ruled out, and all molecular motions along the MTs must have physiological geometry. These end-attachments clearly relied on Ndc80–MT binding moieties because two Ndc80 mutants that perturb MT interactions (K166D and Δ1-80)^25^ failed to maintain end-attachment, whereas the Bonsai mutant, which has a shorter stalk but wild-type MT-binding domains^26^, worked well (Fig. S2d). Thus, a combination of only two proteins, CENP-E kinesin and Ndc80, can provide both efficient lateral transport and wall-to-end transition at stabilized MTs.

### Molecular-mechanical model supports the idea that MT end conversion can be achieved by proteins with no intrinsic MT end-binding activity

To increase our confidence in the above conclusions, we used quantitative methods to recapitulate end conversion *in silico*. Specifically, we employed equations of Brownian dynamics to describe interactions between the MT wall and multiple MT-binding molecules with and without motor activity (see Supplementary Note). In the model, these molecules are immobilized on a surface, representing a patch on the microsphere, as in our end-conversion assay (Fig. 2a, S4). They bind and detach from the MT protofilament stochastically and move randomly (Ndc80) or unidirectionally (CENP-E). Because both the walking motors and thermal fluctuations generate force acting on the MT and this force is transmitted to all MT-bound molecules, the velocity of these molecular motions and the unbinding times were modeled as force-dependent (Fig. 2b,c; Table S1). In this model, as in our experiments, CENP-E brings the MT plus-end to the motor-attachment site. After the last motor walks off the end, the MT detaches from the molecular patch (Video 3). Multiple surface-immobilized molecules of Ndc80, on the other hand, drive lateral MT diffusion, with a rate that decreases as more molecules are modeled (Fig. S3a-c). These diffusive MT motions often (for a few Ndc80 molecules, Video 4) or occasionally (with more molecules) bring one of the MT ends to the molecular patch. Because the patch hosts multiple Ndc80 molecules, the probability of complete MT end detachment is very low, even though each Ndc80 molecule binds for only < 0.5 s^15,27^. The MT end inevitably diffuses away from the edge, centering on average at the Ndc80 patch, as expected from random diffusion of the wall-binding proteins.

**Figure 2.**
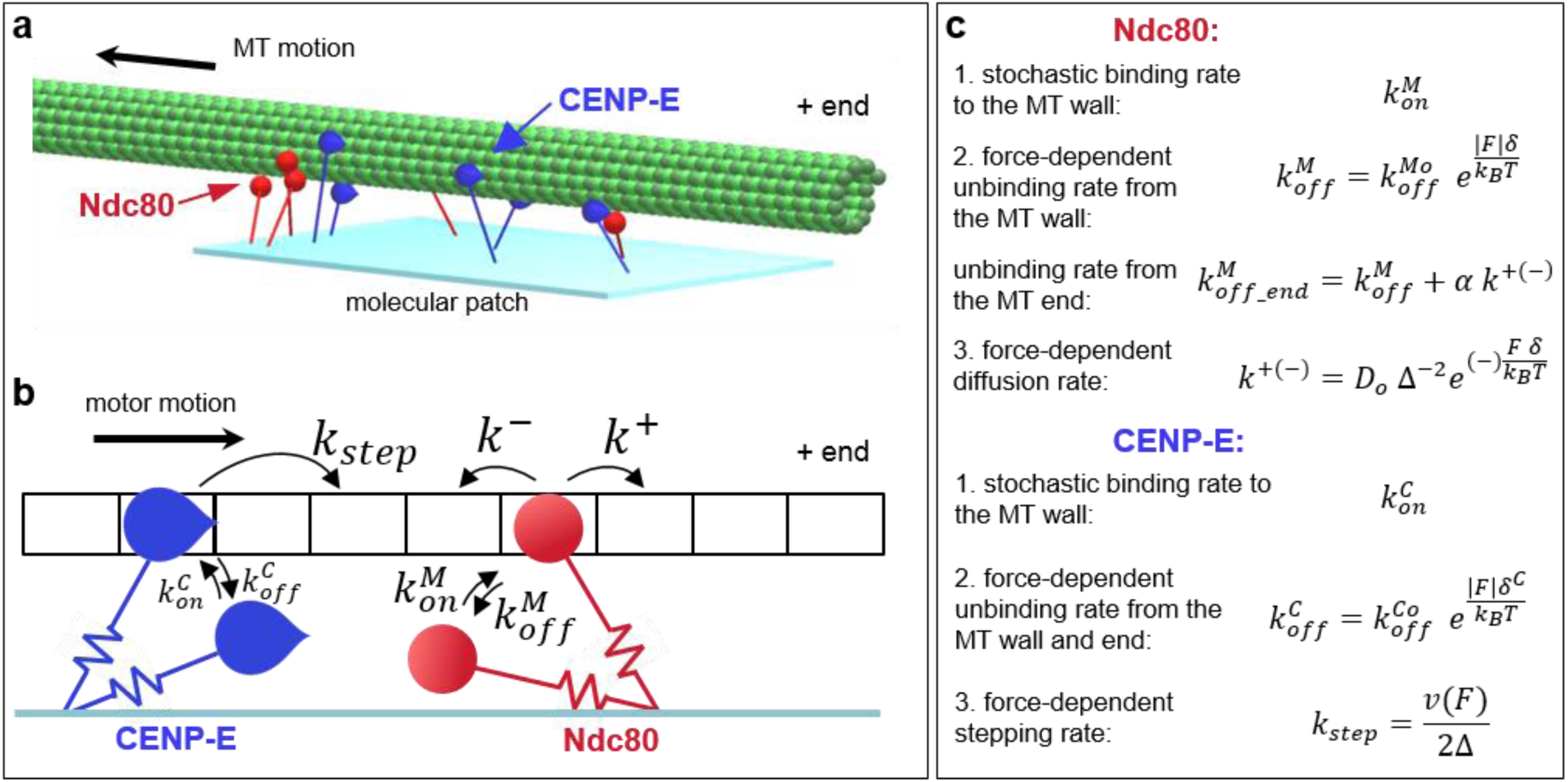
Mathematical model of the molecular ensemble of motors and microtubule-binding proteins moving on the microtubule wall. (a) Multiple MAPs (red) and motors (blue) are randomly distributed on the surface, forming a molecular lawn. Stabilized MT moves under force from kinesin motors in the presence of thermal noise. (b–c) Summary of kinetic transitions. Molecules bind stochastically to the 4-nm sites on the MT wall, and their unbinding is faster under force. Stepping of the motors and diffusional steps of the MAPs are also force-dependent. The motor dissociates from the MT end and the MT wall at the same rate. The MAP molecule can dissociate from the MT end fully or continue to diffuse on the MT wall.

Combining the CENP-E motors and Ndc80 molecules in these simulations revealed highly dynamic and complex ensemble behavior (Video 5). Importantly, the MT only rarely slides all the way to the last motor at the edge of the patch, because as the number of MT-bound motors decreases, they begin to struggle with the MT-bound Ndc80 molecules and frequently dissociate. The MT end, however, does not detach and is even slightly pulled away from the edge by the diffusing Ndc80 molecules. If, however, some or all Ndc80s unbind, the motors resume their persistent transport, trying to decrease the overlap between the MT and the patch. This, in turn, decreases the number of bound motors, the Ndc80/motor ratio increases, and the MT slows down, and the cycle begins again. Because multiple molecules are involved, and their stepping and thermal MT fluctuations are stochastic, these phases of the tip motions are highly irregular, and different numbers of molecules are involved at any given time. Importantly, this complex molecular ensemble can maintain small MT overlap (20–40 nm) for a significant time (>70% of patch–MT attachments lasted >30 min), recapitulating our findings *in vitro*.

### The reconstituted MT end-attachment maintains multiple cycles of MT dynamics

Next, we investigated whether such MT end-attachments could be transformed into dynamic end-coupling. First, we observed fluorescently labeled GMPCPP-stabilized MT seeds gliding along beads coated with Ndc80 and CENP-E, as in the assay with stabilized MTs. Then unlabeled soluble tubulin was added, so that the motions of the labeled MT segments could be seen clearly (Fig. 3a,b). About half of these segments moved slowly away from the beads, indicating that unlabeled tubulin was incorporated at the bead-bound plus ends (Video 6). After several minutes, 83% of these segments rapidly moved back, leading to the saw-toothed distance vs. time plots that are typical for dynamic MTs (Fig. 3c, and 5). The velocity of “away” motion (0.67 ± 0.04 μm/min) was similar to the rate of polymerization of the bead-free MT ends (Fig. 3d). This is significantly slower than CENP-E-dependent MT gliding, suggesting that the “away” motion did not result from lateral CENP-E-dependent gliding. We varied the concentration of soluble tubulin or MgCl_2_, a known effector of MT polymerization^28^. The velocity changed along with the rate of free MT end polymerization (Fig. S5b, Video 7), strongly implying that tubulin assembly was indeed the driving force.

**Figure 3.**
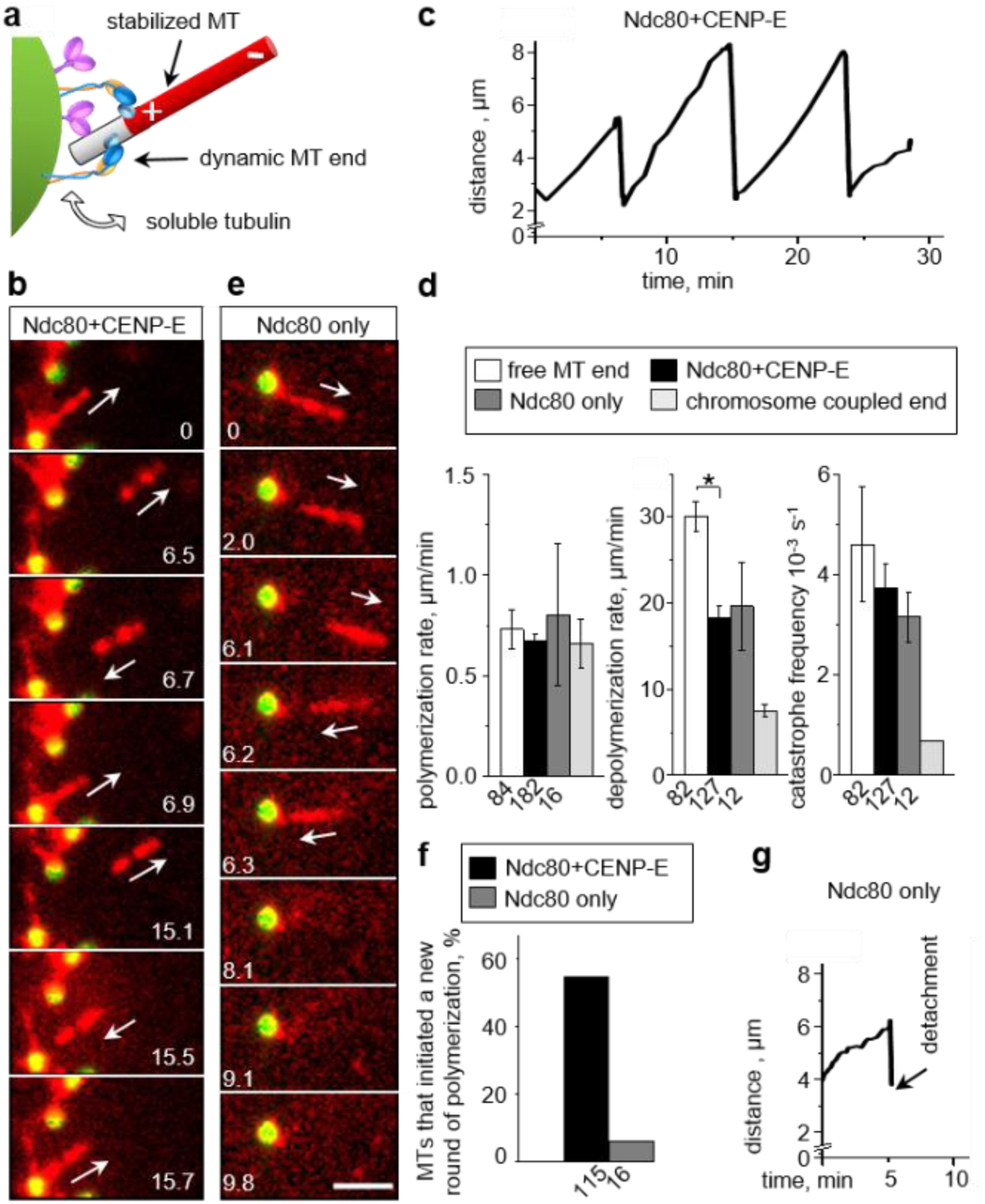
CENP-E and Ndc80 coupling to the dynamic MT ends. (a) Schematics of the dynamic MT end-conversion assay. Gliding of fluorescently-labeled GMPCPP-stabilized MT seeds on beads is observed as in Fig. 1e, and unlabeled soluble tubulin is added to examine its incorporation at the bead-bound MT plus-end. (b) Selected time-lapse images recorded with Ndc80+CENP-E beads after addition of unlabeled soluble tubulin (6.3 µM). Numbers are time (min) from the start of observation. Arrows show the direction of motion of the bright MT fragment, reporting on the dynamics of the bead-bound MT plus-end. Bar, 3 µm. (c) Distance from the distal tip of the fluorescent MT fragment to the bead vs. time, showing repeated cycles. (d) Dynamics parameters for freely growing MTs (*N*=4 trials) and for MT ends coupled to protein-coated beads (*N*=4 for Ndc80+CENP-E; *N*=6 for Ndc80 alone), showing means ± SEM for average results from these trials. Data for isolated mammalian chromosomes are from ref. 30. Statistical differences were evaluated by Kruskal–Wallis ANOVA; * p < 0.05. (e) Images as in panel *b* but recorded using beads coated with Ndc80 only. (f) Percent of bead-coupled MT ends that disassembled, and then initiated a new round of MT polymerization without losing their bead attachment. (g) Plot similar to panel *c* but for a bead coated with Ndc80 in the absence of CENP-E motor. The Ndc80-coated beads can maintain coupling during only one dynamic MT cycle.

To directly determine whether CENP-E was required during the tubulin assembly phase, we examined the dynamics of the bead-coupled MT end in this motor’s absence. A commonly used inhibitor of CENP-E, GSK-923294, induces strong motor-MT binding^29^; consequently, we could not use it in our assay because it would firmly attach MTs to the CENP-E beads. Instead, we took advantage of rare events of direct MT end-binding by chance encounters to beads coated with Ndc80 alone. About 3% of MT segments (n=533) attached via their ends and moved away from the beads after soluble tubulin was added (Fig. 3e). This velocity was very similar to that observed on beads coated with both Ndc80 and CENP-E, consistent with our interpretation that it was determined by tubulin assembly (Fig. 3d). Strikingly, most of the MT segments that moved back to the Ndc80-coated beads after MT catastrophe failed to initiate new “away” motion (Fig. 3f,g and S5c). Thus, CENP-E motor is required for re-establishing dynamic coupling after each depolymerization phase, explaining why the total duration of dynamic coupling was significantly reduced in its absence (Fig. S5c).

Motion of the labeled MT fragment toward Ndc80+CENP-E beads was much faster (18.3±1.3 μm/min) than the “away” motion. Because our assay lacks any depolymerizing or minus-end-directed motors, it could only be driven by MT depolymerization. Consistently, this velocity, like the catastrophe frequency, was not significantly changed in the absence of CENP-E (Fig. 3d). Interestingly, depolymerization of the bead-coupled MT ends was slower than that of the free MT end. This retardation and the slightly reduced catastrophe frequency are consistent with the previously reported suppression of the dynamics of MT ends coupled to the kinetochores of purified mammalian chromosomes *in vitro*^30^. Thus, the MT wall-binding proteins CENP-E and Ndc80 can achieve dynamic end-coupling that is similar to physiological.

### Various MAPs can substitute for Ndc80, but the resultant end-attachments are not as lasting

We applied these experimental and modeling tools to investigate the molecular requirements for end conversion *in vitro*. The essence of our model for the CENP-E–dependent MT wall-to-end transition is that the motor positions Ndc80 molecules near the MT plus-end, where they provide dynamic adhesion via MT wall-diffusion. If this model is accurate, other diffusing MAPs could substitute for the Ndc80 function. To test this prediction, we used four proteins that can diffuse on the MT wall: Ska1 complex^12^, the MT-binding tail of CENP-E^19^ (CENP-E Tail), EB1^31,32^, and CLASP2^33^. Purified GFP-tagged versions of each of these proteins were combined with CENP-E motor (Fig. S6a). CENP-E transported stabilized MTs quickly over these beads, implying that these MAPs imposed less frictional resistance than Ndc80 (Fig. 4a,b). With all these MAPs, we observed that a large fraction of MT plus-ends reached the protein-coated beads and stayed attached for at least 4 s, indicating that these molecular pairs are capable of wall-to-end transition. However, the average duration of end-attachments differed among the MAPs tested: Ska1 and CENP-E Tail maintained end-attachments for only 3–4 min (Video 8), whereas with EB1 and CLASP2, >80% of end-attachments lasted 11-14 min (Fig. 4b,c).

**Figure 4.**
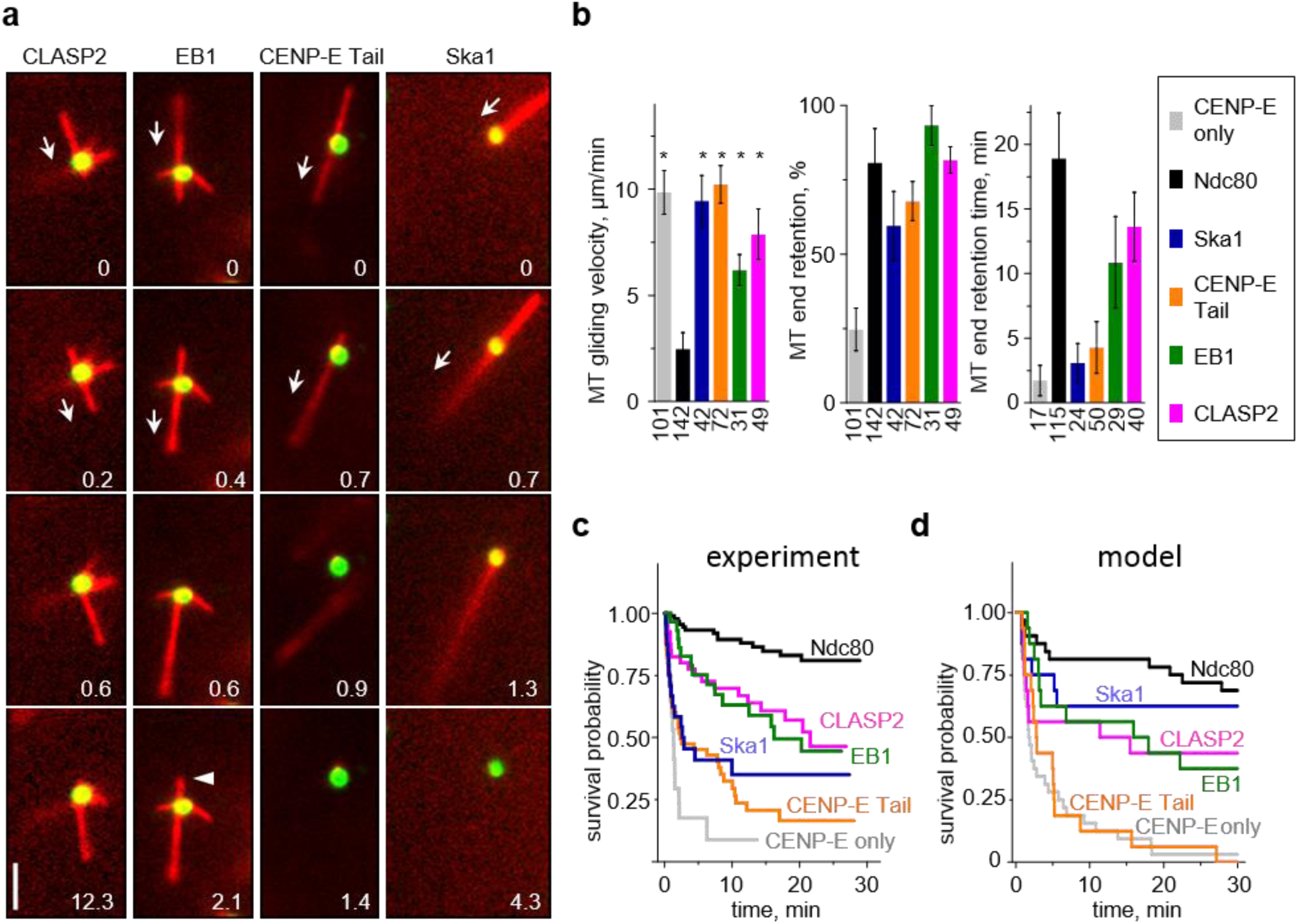
MT wall-to-end transition in molecular systems combining CENP-E motor with various MAPs. (a) Selected time-lapse images of stabilized MTs moving over beads coated with CENP-E motor and the indicated MAP. All proteins were conjugated to beads via anti-GFP antibodies to achieve similar brightnesses, ensuring that any differences in MT interactions are not due to differences in the density of the protein coatings. Numbers are time (min). Arrows show the direction of MT gliding. Bar, 3 µm. Arrowhead in the last EB1 panel points to a loss of tip attachment due to the MT end-to-wall transition. (b) Quantifications as in Fig. 1g, but for beads coated with mixtures containing the CENP-E motor and the indicated MAP. Each MAP was examined in *N*≥3 independent trials, yielding *n* observed events, as indicated below each column. Data are means ± SEM for average results from each trial. Asterisk above a bar (p < 0.05) indicates a significant difference relative to the analogous measurement for Ndc80 beads, as determined by Kruskal–Wallis ANOVA. (c) Kaplan–Meier plot for MT end-retention time based on the same data sets as in panel (b). (d) Kaplan–Meier plot for the predicted MT end-retention time for different MAPs paired with CENP-E motor (*n*=32 simulations for each condition). Different MAPs were modeled using the diffusion coefficients and residency time measured for single molecules *in vitro* (Supplementary Table 2).

To identify the determinants of these differences in end retention, we asked whether our mathematical model could recapitulate these findings based on the quantitative characteristics of these MAPs, such as the rate of MAP diffusion and the duration of one such interaction (Fig. 2c). We used single-molecule visualizations *in vitro* to measure these parameters for Ndc80, Ska1, and CLASP2, and used published results for EB1 and CENP-E Tail (Fig. S3e-g; Table S2). Based on these input parameters, the model correctly describes that Ndc80 has the longest attachment time, CLASP2, EB1, and Ska1 exhibit intermediate end retention and CENP-E Tail has the shortest (Fig. 4d).

Another consistency between our model and experiment is the propensity of MTs to diffuse rapidly on EB1-coated beads. Although EB1 supported wall-to-end transition, the resulting end-attachments *in vitro* were often interrupted by the reverse end-to-wall transitions (29% from n=31 coupled MTs, Fig. 4a arrowhead). During these events, the MT end lost its bead association and diffused vigorously on the bead (Video 9). This behavior is consistent with the fast diffusion rate and short residence time for EB1 (Table S2), which mediates MT diffusion when all CENP-E motors accidentally unbind. By contrast, we did not observe such reverse end-to-wall transitions with Ndc80, as explained in the model by this protein’s relatively slow diffusion and longer residency time (Fig. S5e).

Similar reversions we also observed at dynamic MT ends coupled to the EB1+CENP-E beads (Fig. 5). There were repeated wall-to-end and end-to-wall transitions interspersed with directed transport, so the total end-retention time for EB1+CENP-E beads is likely to be overestimated (Fig. 4b,c). Consistent with our model, no such reversed transitions were observed with CLASP2+CENP-E beads, because CLASP2 diffuses significantly more slowly. Interestingly, CLASP2 could support the “away” motions in the presence of soluble tubulin, but the backward motions were not observed (Fig. 5). Thus, although CLASP2 mediates moderately long attachments of stabilized MT ends, unlike Ndc80 it fails to couple to the ends of depolymerizing MTs.

**Figure 5.**
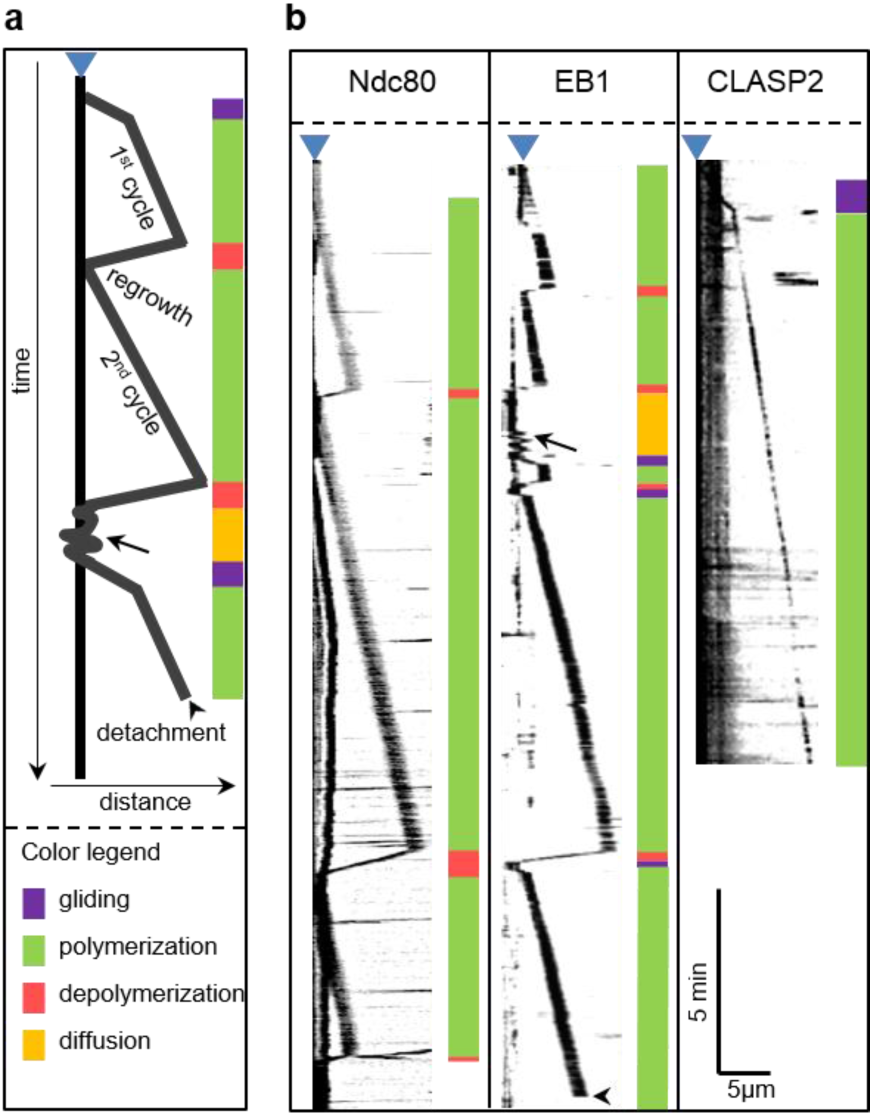
Kymographs from bead coupled MT dynamics experiment. (a) Schematics depicting different features of experimental kymographs with bead-coupled dynamic MT ends. Vertical black line corresponds to a coverslip-immobilized bead (also denoted by blue triangle above), which is often visible in the MT channel owing to the bead-attached motionless MTs. When such MTs were lacking, bead position was determined from GFP channel. The oblique black lines correspond to motions of the brightly labeled GMPCPP-stabilized MT seeds. Color bars on the right provide visual guides for interpretations of these motions. Arrow and arrowhead represent detachment and end to wall transition events respectively. (b) Example kymographs for dynamic MT ends coupled to beads coated with indicated proteins together with CENP-E. See color-coded bars on the right of each kymograph for interpretations.

### At physiological ATP concentration Kinesin-1 cannot replace CENP-E in supporting Ndc80-dependent MT end attachment

To further investigate biophysical mechanisms of end conversion we tried to replace CENP-E with another transporting plus-end-directed motor, Kinesin-1. In the absence of Ndc80, stabilized MTs glided much faster on the Kinesin-1 beads than on beads coated with CENP-E (Fig. 6a,b), consistent with the higher velocity of Kinesin-1^34^. Upon addition of Ndc80, the MT gliding slowed down, and the fraction of MT plus-ends that paused at the beads for > 4 s also increased. However, most of these ends detached quickly at the same Ndc80 concentrations that provided durable attachments to the ends delivered by CENP-E (Video 10). The distinct behaviors of these motors in the MT wall-to-end transition assay was further revealed by plotting the end-retention times of individual MTs against their preceding gliding velocities: although these data were highly variable, the points for CENP-E and Kinesin-1 exhibited minimal overlap (Fig. S6b). Strikingly, MT end retention time by Ndc80 in the presence of Kinesin-1 was much lower than end retention by Ndc80 alone (Fig. S5c). Thus, Kinesin-1 can efficiently deliver the MT plus-end to Ndc80, but it subsequently actively interferes with Ndc80-mediated MT tip attachment.

**Figure 6.**
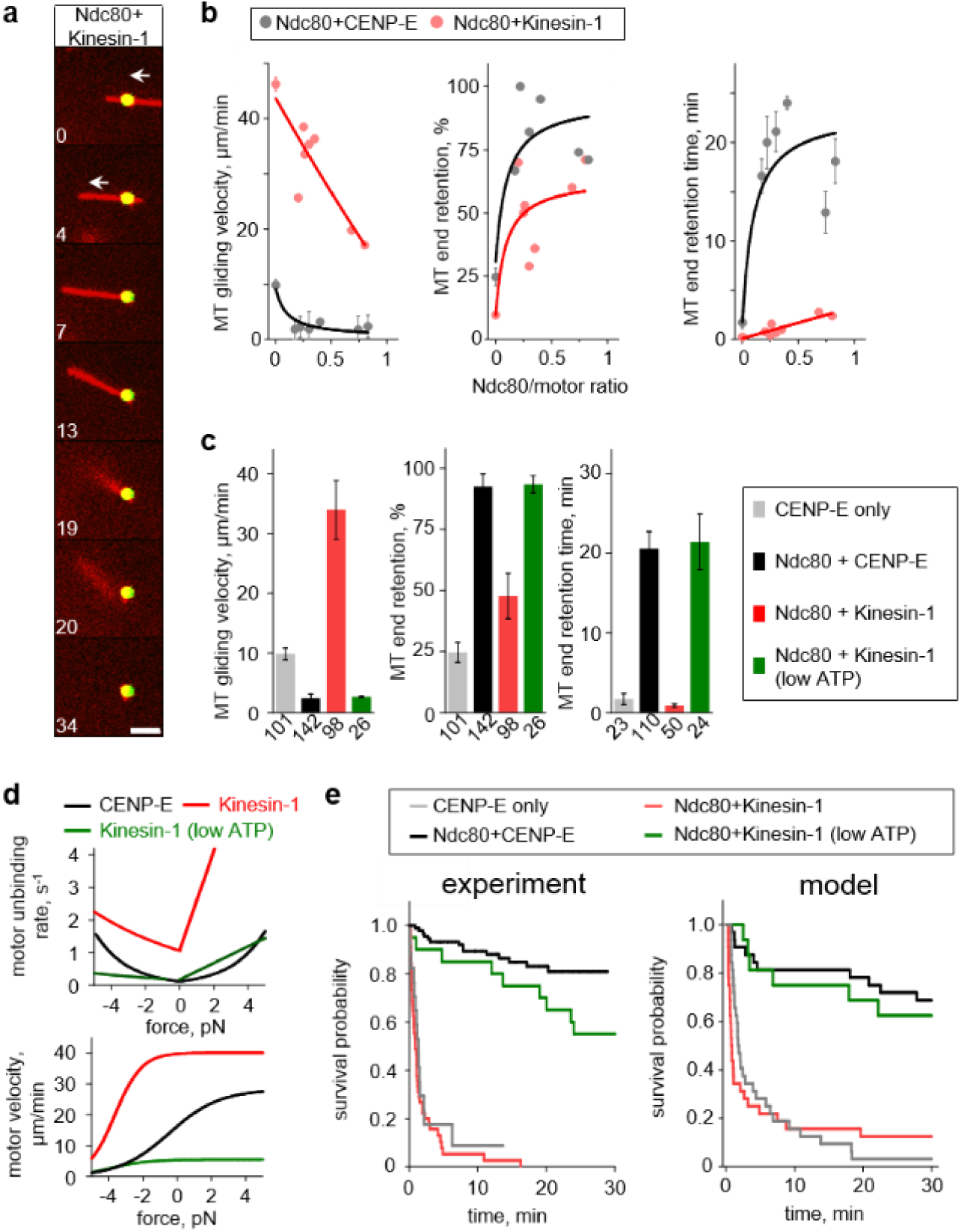
Kinesin-1 paired with Ndc80 in the MT wall-to-end transition assay. (a) Selected time-lapse images of a stabilized MT moving over a bead coated with Kinesin-1 and Ndc80 proteins. Numbers are time (s). Arrows show direction of MT gliding. Bar, 3 µm. (b) MT behavior as a function of the ratio of bead-bound proteins. Points with error bars are means ± SEM calculated for all MTs observed during each experiment. Curves are exponents (left) and hyperbolic functions illustrating the trends. (c) Average results for beads coated with Ndc80 and motors at ratios from 0.3 to 0.5. Data are means ± SEM for the averaged results from each experiment. Numbers under each bar indicate the total number of observed events from at least three experiments. All experiments with motors were carried out in motility buffer supplemented with 2 mM Mg-ATP; low ATP is 20 µM. (d) Force-detachment (top) and force-velocity (bottom) characteristics used in model simulations. Data are based on^18,41^; see Supplement for details. (e) Kaplan–Meier survival plots for the MT end-retention time. Experimental results are as in Fig. 1h but supplemented with measurements for Ndc80+Kinesin-1 at 2 mM ATP (*N*=7, *n*=50) and 20 µM ATP (*N*=3, *n*=24). Theoretical plot is based on *n*=32 simulations for each condition modeled using characteristics in panel *d*.

To obtain mechanistic insight into this unexpected result, we turned to our mathematical model, in which motor function is described using two force-dependent characteristics, for velocity and unbinding (Fig. 2c). Although CENP-E is thought of as a mitotic version of Kinesin-1^35^, the motility characteristics of the two proteins under force are markedly different (Fig. 6d)^20,35,36^. When these Kinesin-1 characteristics were incorporated, the model correctly predicted a low survival probability for MT-bead attachments (Fig. 6e). We used this tool to find conditions that “convert” Kinesin-1 into CENP-E, and discovered that end retention could be extended *in silico* using Kinesin-1 characteristics corresponding to 100-fold lower ATP concentration, which decreases the unbinding rate and motor’s velocity^37^ (Fig. 6d; Supplementary Note). We tested this prediction experimentally. At 20 µM ATP, the MT gliding velocity on Ndc80+Kinesin-1 beads decreased (Fig. 6c), as expected. Consistent with the model prediction, the arriving MT ends remained attached to these beads for much longer (Fig. 6e). To uncouple the impact of velocity retardation from more stable Kinesin-1 MT association we modeled these conditions separately; the longer end-retention time could be obtained using the modified Kinesin-1 force-unbinding characteristic and its naturally high velocity but not vice versa (Fig. S4c). Thus, the distinct end-retention behavior of CENP-E and Kinesin-1 motors stems from their different unbinding rates under force.

## DISCUSSION

MT end conversion by mitotic kinetochores is one of the least understood transitions that occurs during chromosome segregation. Until now it has not been reconstructed and analyzed using quantitative approaches. Previous experiments using Ndc80-coated beads mimicked MT end conversion by triggering depolymerization of the laterally attached MT, causing the bead to follow the depolymerizing tip^17^. Other studies used laser beams to promote direct binding between MT tips and Ndc80 beads, which subsequently remained attached to the dynamic ends for few minutes^13^. It remains unclear whether these events reflect natural end conversion, especially because MAP-coated beads can follow MT tips while rolling^38^. The approach described here recreates a more physiologically relevant situation by introducing the motor domains of the plus-end–directed CENP-E kinesin, which during chromosome congression transports the Ndc80-containing kinetochores.

Our novel assay reveals that the CENP-E motor and Ndc80 complex represent an optimally tuned molecular system capable of MT end conversion *in vitro*. During lateral transport, Ndc80 binding to the MT wall creates drag that antagonizes the walking of the CENP-E motor, suggesting that the 10-fold lower velocity of chromosome congression relative to freely walking CENP-E^39^is a result of molecular friction from kinetochore-bound Ndc80 complexes. Near the MT plus-end, CENP-E and Ndc80 generate a physiologically competent MT attachment, as seen from its ability to persist during several dynamic cycles of the coupled MT end, lasting tens of minutes.

Several lines of evidence suggest that the underlying molecular interactions occur at the wall of the MT near its tip, rather than requiring these proteins to bind specifically to the protruding protofilaments, which are more curved than protofilaments within intact MT walls^17,40^. First, both CENP-E motors and Ndc80 are MT wall-binders with no intrinsic ability to bind MT tips *in vitro*^12,16,19^. Second, these molecules are immobilized randomly on the bead surface, rather than clustered in one spot; therefore, the most likely configuration is lateral attachment of these molecules along the MT wall segment immediately adjacent to the tip, as seen in our model Video 5. Third, end conversion is not a unique property of these two proteins. Other MAPs can substitute for Ndc80, but there is no clear correlation between their performance in this assay and their MT tip-binding properties. For example, EB1 strongly prefers growing, but not depolymerizing, MT tips^31^ whereas the Ska1 complex can track both growing and shortening MT ends^12,41^. Both proteins, however, were less effective than Ndc80 in the end-conversion assay (Fig. 4c, S6c).

Interestingly, the duration of MT end retention correlates with the velocity at which CENP-E transports MTs laterally bound to these MAPs (Fig. S6c), suggesting that maintenance of end attachment is determined mostly by the MT wall-binding properties of these MAPs. Indeed, the survival plots for the MT end-retention time for these MAPs were similar to those obtained in the model using only two input parameters, the diffusion rate and residence time (Fig. 4c,d), both of which describe MT wall-binding. According to the model, Ndc80 provides the best end retention among the tested MAPs due to a combination of slow diffusion and relatively long residence time. Interestingly, the retardation of CENP-E transport by Ndc80 predicted by the model was not as strong as that observed in experimentally. This suggests that Ndc80 motility under force may differ from that of other MAPs, an important prediction that should be tested in future work.

Another inconsistency between theory and experiment is that Ska1 protein has a shorter end-retention time *in vitro* than predicted by the model. This could be explained by some aspect of the experimental system, such as suboptimal activity of bead-immobilized Ska1 due to the specific location of the tag. Alternatively, this discrepancy could indicate that MT-binding characteristics that are not yet included in our model, or are assumed to be equivalent for all MAPs, also influence end-retention time. There is impressive quantitative consistency between our experimental observations and the model’s prediction of end conversion with Kinesin-1, suggesting that our interpretations are mechanistically accurate. This kinesin proficiently delivers the MT plus-ends to the bead-conjugated Ndc80 molecules, but in contrast to the situation in the presence of CENP-E, the Ndc80 molecules fail to provide lasting attachment to these ends. Reducing the ATP concentration dramatically increases the end-retention time, revealing an essential role for the force-sensitivity of motor walking on the MT.

Thus, the motility characteristics of the motors and MAPs forming a multimolecular ensemble must be finely tuned to enable emergent end-retention behavior. These molecules maintain a short (20–40 nm) overlap with the distal MT wall segment, similar to the overlap created by the multimolecular “sleeve” in Hill’s model^42^, although the underlying biophysical mechanisms are different^10^ Because the stepping of the MAPs forming the sleeve is highly coordinated, it tends to engage maximal number of MAPs, corresponding to the minimal free energy of this system. In our model, all molecular interactions are uncoordinated and stochastic, as proposed in the kinetochore “molecular lawn” representation^43^. Here, entropic forces play a significant role, and the number of MT-bound molecules is limited by kinetic binding constants^44^. These factors, together with the force-sensitivity of stepping and unbinding, determine the outcome of end retention.

Our model lays a novel conceptual foundation for a quantitative molecular understanding of chromosome motility during mitosis. The proposed mechanism explains CENP-E–mediated end conversion without invoking specialized tip-binding proteins or regulatory modifications (which may nonetheless provide additional layers of complexity to kinetochore–MT interactions in cells). The end-on MT attachments in metaphase cells have a markedly different geometry than those in our reconstitutions. We hypothesize that end-on configuration in cells results from two distinct factors: molecular MT-wall interactions and mechanical forces. In this view, the initial MT-lateral attachment of the kinetochores is first replaced by binding to the terminal MT-end segment lying obliquely to the molecular lawn on the kinetochore surface (Fig. 7a,b). The MT-perpendicular end-on configuration is then induced by spindle forces that orient sister kinetochores along the spindle axis (Fig. 7c). We propose that despite these different MT-kinetochore orientations, the nature of the underlying molecular interactions remains the same: in the MT-perpendicular end-on configuration, Ndc80 and CENP-E continue to interact with the tip-adjacent MT wall via essentially same biophysical mechanism as in the oblique configuration (Fig. 7c). Thus, this model provides a consistent and unifying molecular basis for the initial lateral transport, wall-to-end transition, and durable end-coupling at vertebrate kinetochores.

**Figure 7.**
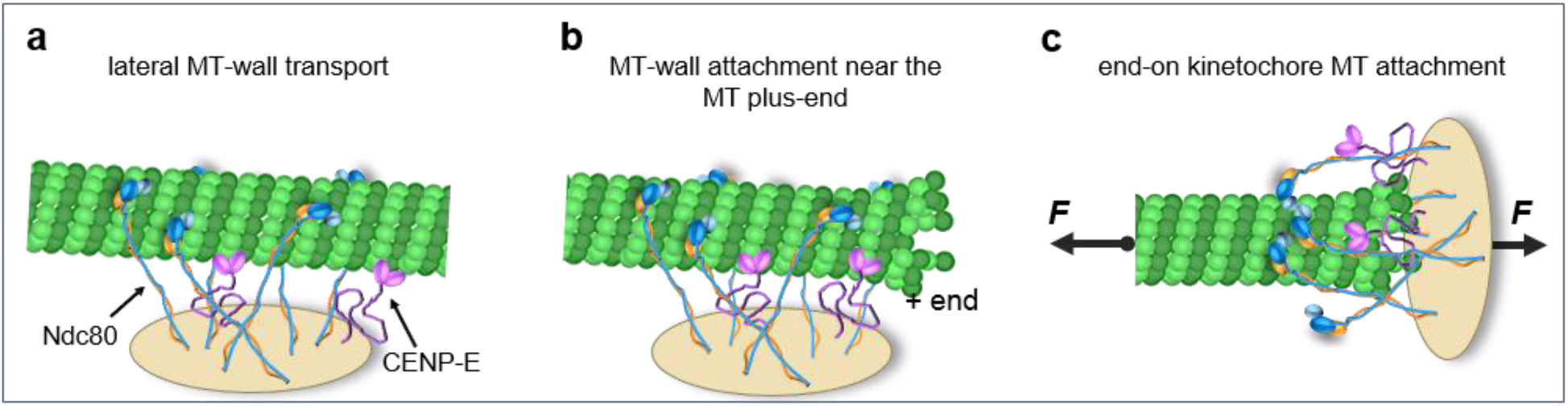
Molecular and MT configurations during wall-to-end transition. Multimolecular ensemble of CENP-E kinesins and Ndc80 complexes, forming a molecular lawn that interacts with the MT wall. Ndc80 slows down CENP-E kinesin during the plus-end–directed transport (a), but plays an essential role in attachment to the end-proximal MT wall. In our *in vitro* experiments, the molecular interactions are concentrated at one side of the MT, which forms oblique contact with the bead surface, as shown by *in silico* simulations (b). This configuration is also likely to occur transiently at the kinetochores of mitotic cells. However, forces acting on the chromosomes and kinetochore-bound MTs reorient the kinetochore, promoting the classical end-on configuration (c). We propose that this attachment is mediated by essentially the same molecular interactions with the MT wall, as described in this work.

## Supporting information

## ACKNOWLEDGEMENTS

Plasmids and protein purification protocols were generously provided by Drs. I. Cheeseman (Whitehead Institute, MIT), J. DeLuca (Univ. of CO at Ft. Collins), T. Surrey (Francis Crick Inst., UK), and D. W. Cleveland (Ludwig Institute for Cancer Research and Univ. of CA at San Diego). We are grateful to Drs. H. Maiato and S. Macedo-Ribeiro (Instituto de Investigação e Inovação em Saúde, Universidade do Porto, Portugal) for providing CLASP2 protein. We also thank Dr. A. Kiyatkin, P.-T. Chen and V. Mustyatsa for help with protein purification, and Grishchuk lab members for discussions. Funding for this work was provided by the National Institutes of Health grant GM-R01098389 to E.L.G., and American Cancer Society grant RSG-14-018-01-CCG to E.L.G. A.C.F. acknowledges the support by CODECHECK grant from the European Research Council, FLAD Life Science 2020, and the Louis-Jeantet Young Investigator Career Award (to H. Maiato). F.I.A. acknowledges support from the Russian Foundation for Basic Research 15-04-04467 and 16-34-60196 MOL_A_DK.

## AUTHOR CONTRIBUTIONS

M.C., E.T. and A.V.Z. performed experiments, A.V.Z., M.G. and F.I.A. carried out mathematical modeling, A.C.F. purified human CLASP2 protein, E.L.G and M.C. designed research, analyzed data and wrote the paper with input from A.V.Z, E.T. and F.I.A.

## METHODS

### Protein purification

Tubulin from bovine brain was purified by thermal cycling and chromatography and labeled with HiLyte647 as in^45^. The *Xenopus laevis* CENP-E-GFP construct (1-473 aa) was expressed and purified from *Escherichia coli* as in^46^. This truncated CENP-E protein contains the motor domains of CENP-E dimerized with a short segment of native stalk, but the rest of the stalk and the MT-binding tails are absent (Fig. S1a). Truncated CENP-E with C-terminal Myc-tag in place of GFP was a gift from Drs. Y. Kim and D. W. Cleveland. The GFP-tagged C-terminus of CENP-E (CENP-E Tail) was purified as in^19^; human Bonsai Ndc80-GFP protein complex as in^25^; human Broccoli Ndc80-GFP construct as in^12^; full length human EB1-GFP as in^47^; GFP tagged Kinesin-1 (1–560 aa) as in^21^; and full length human Ska1-GFP complex consisting of all three subunits Ska1, Ska2, and Ska3 as in^48^. Human GFP-CLASP2 was purified using Baculovirus Expression Vector System (BEVS) (BD Biosciences, San Jose, CA, USA). Briefly, CLASP2 ORF was subcloned into expression vector pKL in fusion with an N-terminal GFP-tag and a C-terminal His10-tag. The resultant plasmid, pKL-GFP-CLASP2-His10x, was used to transform DH10EmBacY *E. coli* for transposition into bacmid. Production of recombinant baculovirus and transfection of *Spodoptera frugiperda* Sf21 cells was performed using the MultiBac expression system^49^. Sf21 cells were lysed by sonication in 50 mM Tris-HCl pH 8.0, 150 mM NaCl, 7 mM β-mercaptoethanol supplemented with protease inhibitors (Complete EDTA-free, Roche, Basel, Switzerland). Clarified protein extracts were loaded onto a HisTrap HP column (GE Healthcare, Chicago, IL, USA) pre-equilibrated in 50 mM Tris-HCl pH 8.0, 500 mM NaCl, 20 mM imidazole, 7 mM β-mercaptoethanol and eluted with 200 mM imidazole. CLASP2-containing fractions were pooled and further purified on a HiPrep 16/60 Sephacryl S-300 HR column (GE Healthcare) pre-equilibrated with 50 mM sodium phosphate buffer pH 7.0, 400 mM KCl, 2 mM MgCl_2_, 0.8 mM β-mercaptoethanol, 1% glycerol.

### Fluorescence microscopy assays

Prior to each motility experiment, a frozen protein aliquot was thawed and clarified by ultracentrifugation (TLA100 rotor, Beckman Coulter, Brea, CA, USA) at 156,845 × *g* for 15 min at 4°C. Protein concentration in the supernatant was determined by measuring GFP intensity by fluorescence microscopy and comparing to a “standard” GFP-labeled protein whose concentration was determined by spectrometry ^50^. For gliding assays, taxol-stabilized MTs were prepared from unlabeled and HiLyte647-labeled tubulin in a 24:1 ratio. For MT wall-to-end transition assays, stabilized MTs were prepared from a mixture of unlabeled and HiLyte647-labeled tubulin (10:1, total tubulin concentration, 72.5 µM) and 1 mM GMPCPP (Jena Bioscience, Jena Germany) incubated at 37°C for 10 min. All motility assays were carried out using a Nikon Eclipse Ti-E inverted microscope equipped with 1.49x NA 100x oil objective and Andor iXon3 CCD camera (Cambridge Scientific, Watertown, MA, USA), as in ^19^. Under these conditions, the microscope produced 512 × 512-pixel images with 0.14 µm/pixel resolution in both the *x* and *y* directions. All experiments were carried out at 32°C by heating the objective with an objective heater (Bioptechs, Butler, PA, USA).

### Motility assays using free-floating beads and a laser trap

Assays were carried out as in^20^. COOH-activated glass beads (0.5 μm, Bangs Laboratories, Fishers, IN, USA) were coated with a mixture of Broccoli Ndc80-GFP and truncated CENP-E-GFP using 12-nm DNA links as in^20^. The ratio of these proteins was varied while holding constant the combined total protein concentration (40 nM). In experiments with dynamic MTs, the Ndc80/CENP-E ratio was 0.25. Because both proteins were conjugated to beads through a GFP-tag, this ratio reflects the proportion of Ndc80 and CENP-E molecules conjugated to the surface of the beads. Perfusion chambers were prepared by attaching a silanized glass coverslip over a regular glass slide with double-sided sticky tape (Scotch) to generate a 15-µl flow chamber as in^51^. Solutions were exchanged using a peristaltic pump as in^51^. Taxol-stabilized MTs were prepared and immobilized on the coverslip using anti-tubulin antibodies (Serotec) as in^20^. For experiments with dynamic MTs, digoxigenin (DIG)-labeled GMPCPP-stabilized MT seeds were prepared and immobilized on the coverslip via anti-DIG antibodies (Roche) as in^41^. Next, beads were flowed into the chamber in “imaging” buffer: BRB80 (80 mM PIPES, 4 mM MgCl_2_, 1 mM EGTA, pH 6.9) supplemented with 4 mg/ml BSA, 2 mM DTT, 2 mM Mg-ATP, 6 mg/ml glucose, 80 μg/ml catalase, and 0.1 mg/ml glucose oxidase. For experiments with taxol-stabilized MTs, this buffer was further supplemented with 0.5% β-mercaptoethanol and 7.5 μM taxol. For experiments with dynamic MTs, the imaging buffer was supplemented with 6 μM soluble tubulin and 1 mM Mg-GTP. Using the 1064-nm laser beam of our laser trap^34^, a free-floating bead was trapped and brought into contact with the wall of the taxol-stabilized MT^20^, examining 4-12 beads for each Ndc80/CENP-E ratio. In experiments with dynamic MTs, the beads attached spontaneously to the MTs. Image acquisition in differential interference contract mode was performed with exposure times of 100 ms using either a Cascade 650 CCD (Photometrics, Tucson, AZ, USA) or an Andor iXon3 camera controlled by the Metamorph software (Molecular Devices, San Jose, CA, USA) or NIS-Elements software (Nikon Instruments, Melville, NY, USA) correspondingly.

### MT gliding assay

Perfusion chambers and coverslips coated with proteins were prepared as described in^50^ using a mixture of 0.1 µM biotinylated anti-Myc antibodies (EMD Millipore, Burlington, MA, USA) and 0.1 µM biotinylated anti-GFP antibodies (Abcam, Cambridge, MA, USA) in wash buffer (BRB80 supplemented with 4 mg/ml BSA, 0.1 mM Mg-ATP, and 2 mM DTT). A solution of 0.1 µM Myc-CENP-E in wash buffer was added for 30 min, and then the chamber was washed and incubated with Bonsai Ndc80-GFP at 10–150 nM for 30 min. TIRF (Total Internal Reflection Microscopy) images of five different fields were collected with exposure times of 300-ms exposure using a 488-nm laser. The brightness of Bonsai Ndc80-GFP intensity was calculated by averaging mean intensities of five different fields using the ImageJ software^52^. The resulting density of Bonsai Ndc80-GFP coating, as judged by GFP fluorescence, increased linearly with increasing concentration of soluble protein (Fig. S1b). Then, taxol-stabilized HiLyte647-labeled MTs were introduced in imaging buffer (described in “Motility assays” above) supplemented with 10 µM taxol, the chamber was sealed with VALAP (1:1:1 vaseline:lanolin:paraffin), and gliding MT motions were recorded continuously using TIRF mode for 10 min. MTs with approximately linear trajectories were selected, and their velocities were estimated from kymographs.

### MT wall-to-end transition assay

Perfusion chambers with coverslip-immobilized microbeads were prepared as in^50^. Briefly, streptavidin-coated 0.9-µm polystyrene beads bound to coverslips were coated with biotinylated anti-GFP antibody (0.1 µM), and subsequently blocked with 100 µM biotinylated PEG, resulting in negligible MT binding to bead and coverslip surfaces. A solution of 0.3 µM CENP-E-GFP or Kinesin-1-GFP in wash buffer was added and incubated for 30 min, and several images of beads were collected for subsequent quantifications of GFP intensity, corresponding to the density of motor coating. Images of beads coated with these and other GFP-labeled proteins were acquired with a 488-nm 100-mW diode laser (Coherent, Santa Clara, CA, USA) at 10% power with an exposure time of 300 ms. The chambers were then washed and incubated for 30 min with a GFP-labeled MAP. Unless stated otherwise, the MAP concentrations were as follows: Broccoli Ndc80 0.4 µM; CLASP2 0.4 µM; Ska1 0.2 µM; CENP-E Tail 0.1 µM; EB1 0.4 µM. Images of the beads were collected again to record the increase in GFP intensity. The resultant MAP coatings were similar in density (see Fig. S6a), as judged by the GFP intensities of the beads. Next, GMPCPP-stabilized MT seeds in warm wash buffer in which ATP was replaced with 0.1 mM AMP-PNP (Sigma-Aldrich) were introduced to promote MT binding to the motor molecules on the beads. Then, imaging buffer with 1 mM ATP was washed in and epifluorescence images of HiLyte647-labeled MTs were collected every 4 s for 30 min with exposure times of 300 ms using a 70-mW 640-nm diode laser (Coherent) at 50% power. Videos were prepared by merging channels from MT and bead fields using the Metamorph software (Molecular Devices, San Jose, CA, USA).). In some experiments, AMP-PNP step was omitted, MTs in imaging buffer with 1 mM ATP were added to the protein-coated beads, and the samples were viewed immediately. The results obtained with and without AMP-PNP were very similar (Fig. S2e), so they were combined into a single dataset. For experiments with Kinesin-1 at low ATP concentration, the imaging buffer was supplemented with 20 µM Mg-ATP instead of 2 mM Mg-ATP. MT end-conversion (coupling to dynamic MT ends) was examined with GMPCPP-stabilized MT seeds, except that after these MTs glided for 30 min, unlabeled soluble tubulin (6 µM) in imaging buffer supplemented with 1 mM Mg-GTP was added. This tubulin solution (40 µl) was perfused for 2 min with a peristaltic pump at 20 µl/min, and imaging continued at 60 frames/min for an additional 30 min. Note that in this assay, MT images were collected using epi-fluorescence, rather than TIRF, to increase imaging depth. MTs bind all over the surface of the 1 µm bead, rendering most of bead-bound MTs invisible in TIRF. Because epi-fluorescence illumination has high background when soluble fluorescently-labeled tubulin is used, MT elongation in this assay could only be examined using unlabeled tubulin. Under these conditions, motions of the brightly labeled seeds away and toward the bead could be clearly followed, allowing the unambiguous determination of the site of tubulin addition/loss as the bead-bound MT plus-end.”

### Quantitative data analyses

The brightness of beads coated with different GFP fusion proteins was measured as the integral intensity of the 3.5 × 3.5 µm area encompassing the bead minus the intensity of the same area at a nearby location. MAP-GFP brightness was calculated as the GFP fluorescence intensity of a bead after incubation with the MAP-GFP minus the intensity of the same bead before the MAP-GFP was added; the latter corresponds to the brightness motor-GFP. The ratio of Ndc80/motor (CENP-E or Kinesin-1) brightness was doubled to take into account the fact that each Ndc80 molecule has only one GFP, whereas all other tested MAPs are homodimers.

In the MT wall-to-end transition assays, only MTs that satisfied all of the following selection criteria were selected for quantitative analysis: the MT should be clearly visible and be in focus; the MT should not simultaneously contact several beads, and the MT must have glided until its trailing end reached the bead. The position of the bead in the kymograph was determined by merging the bead and MT images. A successful MT end retention event was counted only if the trailing MT end was coupled to the bead for longer than two successive frames (4 s). The percentage of end-retention events was calculated from the total number of trailing MT ends reaching the beads. Total MT end-retention time was calculated from kymographs as the time between the end of MT gliding motion and the loss of MT attachment or the end of imaging. Due to the latter events, average end-retention times were underestimated. To overcome this limitation, survival probabilities for bead-coupled MT ends were plotted using the Kaplan–Meier algorithm implemented with the Origin software (OriginLab, Northampton, MA, USA).

In experiments containing soluble tubulin, suspected MT polymerization–driven “away” motion was identified when a fluorescent MT seed moved sufficiently far from the bead to leave a visible gap between the fluorescently-labeled MT segment and the bead. The fraction of dynamic MT attachments was calculated as the ratio of the number of MTs that exhibited at least one “away” motion to the total number of MTs bound to beads. Velocities of the away/backward motions, their respective durations, and the maximum lengths of polymerized MTs for each dynamic cycle were estimated from kymographs. The velocities of polymerization and depolymerization motions for free MT ends were calculated in an analogous manner. The total dynamic MT attachment time was calculated as the time between the start and end of all dynamic motions or the end of imaging. MT catastrophe frequency was calculated by dividing the observed number of catastrophes by the sum of the elongation time for all polymerizing MTs.

### Measurement of the MT diffusion on the Ndc80-coated beads

Experimental chambers and solutions were prepared as for the MT wall-to-end transition assay except that beads were coated only with Ndc80-GFP to achieve same Ndc80-GFP brightness as in the wall-to-end transition assay. GMPCPP-stabilized MT seeds were perfused into the chamber and imaged for 30 min at 15 frames/min in appropriate emission channel using same acquisition settings as in end-conversion experiments. Only those bead-bound MTs that satisfy following criteria were selected: MT should be bound only to a single bead; both MT ends should be in focus and clearly visible; during diffusion MT should only diffuse laterally attached to the bead and avoid tip-attachment. One end of each of these MTs was tracked manually using ImageJ to obtain distance vs time track. Next, for each track initial position was subtracted, and squared displacement (MSD) vs time was calculated for each tracked MT. To obtain mean squared displacement plot (Fig. S3b) squared displacements were averaged between all MTs. Diffusion coefficient was determined as a half of the slope of mean squared displacements of these MT ends over time (Fig. S3c).

### Measurement of the dynamics of free MT ends

Dynamics of MT ends that were not bead-bound were examined using the same assay conditions as for MT ends coupled to beads. Coverslip-immobilized streptavidin beads were incubated with 0.1 µM biotinylated anti-mouse IgG antibody (Jackson Immuno Research, West Grove, PA, USA) for 15 min, followed by 0.1 µM mouse anti-tubulin antibody (AbD Serotec/Bio-Rad, Hercules, CA, USA) for 15 min. HiLyte647-labeled GMPCPP-stabilized MT seeds in warm wash buffer were allowed to bind to these beads, and dynamic MT segments were generated by adding a mixture of unlabeled and HiLyte-labeled tubulins (15:1 ratio, total tubulin concentration, 6.3 µM) in imaging buffer supplemented with 1 mM Mg-GTP. Images were recorded in TIRF mode for 30 min with exposure times of 300 ms at 60 frames/min. Data in Fig. 3d for the chromosome-coupled MT end is from ^28^ for a tubulin concentration of 6.8 μM; catastrophe frequency for these ends was calculated as the inverse of the average time until catastrophe (Table 1 in^30^).

### Single-molecule TIRF assay and data analysis

Diffusion of different MAPs on the GMPCPP-stabilized MT seeds labelled with HiLyte-645 was examined as in^27^ using same imaging buffer as in the MT wall-to-end transition assay. Concentrations were selected to achieve single-molecule decoration of MTs and were 0.1 nM for Ndc80, 0.15 nM for Ska1 and 0.3 nM for CLASP2. Exposure times *t*_*exp*_ were 20 ms for Ndc80 and 10 ms for Ska1 and CLASP2. Laser power used was 75% for Ndc80 and 100% for Ska1 and CLASP2. To determine diffusion coefficient for each MAP, MSD was plotted vs. time based on >100 tracks of diffusing molecules (Fig. S3f). Residence time was determined by analyzing all molecular tracks durations. Tracks durations were recorded by clicking on their first and last points using custom-written program in Mathematica (Wolfram Research). Next, in order to correct for undercount of short-lived binding events, we introduced a threshold on a collected residence times – only events longer than 8 *texp* were included to plot the cumulative distribution. These cumulative distributions were then fitted with an truncated exponential distribution:

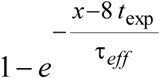

Because the effective residence time is shorter than real molecular residence time due to photobleaching, the kinetic rate constant of GFP photobleaching *k*_*bleach*_ under our typical imaging conditions was determined as in^27^, and the residence time τ was calculated using following expression:

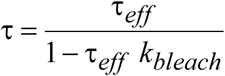

In our experiments *k*_*bleach*_ was 0.17 s^-1^ for Ndc80 complex (at 75% of laser power), and 0.21 s^-1^ for other proteins (measured at 100% laser power) (photobleaching rate was measured as in^27^).

### Theoretical modeling

Mathematical model is done using Brownian dynamics stochastic simulation of ensemble of MAPs and molecular motors stochastically interacting with MT. To describe these stochastic molecular interactions, we used force-dependent characteristics for motors and MAPs stepping and unbinding from the MT. These molecular characteristics for motors and MAPs were based on *in vitro* measured parameters. For complete description of mathematical model see Supplementary Note. Code for simulations was written in Mathematica (Wolfram Research).

## Abbreviations

MAP: microtubule associated protein
MT: microtubule
MSD: mean squared displacement
TIRF: total internal reflection fluorescence

## REFERENCES

1. Walczak, C. E., Cai, S. & Khodjakov, A. Mechanisms of chromosome behaviour during mitosis. Nat. Rev. Mol. Cell Biol. 11, 91–102 (2010).

2. Wood, K. W., Sakowicz, R., Goldstein, L. S. B. & Cleveland, D. W. CENP-E is a plus end-directed kinetochore motor required for metaphase chromosome alignment. Cell 91, 357–366 (1997).

3. Yen, T. J. et al. CENP-E, a novel human centromere-associated protein required for progression from metaphase to anaphase. EMBO J. 10, 1245–1254 (1991).

4. McIntosh, J. R., Grishchuk, E. L. & West, R. R. Chromosome-microtubule interactions during mitosis. Annu. Rev. Cell Dev. Biol. 18, 193–219 (2002).

5. Tanaka, K. Dynamic regulation of kinetochore-microtubule interaction during mitosis. J. Biochem. 152, 415–424 (2012).

6. Sikirzhytski, V. et al. Microtubules assemble near most kinetochores during early prometaphase in human cells. J Cell Biol. 217, 2647–2659 (2018).

7. Rieder, C. L. & Salmon, E.. The vertebrate cell kinetochore and its roles during mitosis. Trends Cell Biol. 8, 310–318 (1998).

8. Shrestha, R. L. & Draviam, V. M. Lateral to end-on conversion of chromosome-microtubule attachment requires kinesins cenp-e and MCAK. Curr. Biol. 23, 1514–1526 (2013).

9. Godek, K. M., Kabeche, L. & Compton, D. A. Regulation of kinetochore-microtubule attachments through homeostatic control during mitosis. Nat. Rev. Mol. Cell Biol. 16, 57–64 (2015).

10. Grishchuk, E. L. Biophysics of Microtubule End Coupling at the Kinetochore. Prog. Mol. Subcell. Biol. 56, 397-428 (2017)

11. Cheeseman, I. M. & Desai, A. Molecular architecture of the kinetochore-microtubule interface. Nat. Rev Mol. Cell Biol. 9, 33–46 (2008).

12. Schmidt, J. C. et al. The kinetochore-bound Ska1 complex tracks depolymerizing microtubules and binds to curved protofilaments. Dev. Cell 23, 968–80 (2012).

13. Powers, A. F. et al. The Ndc80 kinetochore complex forms load-bearing attachments to dynamic microtubule tips via biased diffusion. Cell 136, 865–875 (2009).

14. Lampert, F., Hornung, P. & Westermann, S. The Dam1 complex confers microtubule plus end-tracking activity to the Ndc80 kinetochore complex. J. Cell Biol. 189, 641–649 (2010).

15. Umbreit, N. T. et al. The Ndc80 kinetochore complex directly modulates microtubule dynamics. Proc. Natl. Acad. Sci. U. S. A. 109, 16113–8 (2012).

16. Alushin, G. M. et al. The Ndc80 kinetochore complex forms oligomeric arrays along microtubules. Nature 467, 805–10 (2010).

17. McIntosh, J. R. et al. Fibrils connect microtubule tips with kinetochores: a mechanism to couple tubulin dynamics to chromosome motion. Cell 135, 322–33 (2008).

18. McIntosh, J. R., Volkov, V., Ataullakhanov, F. I. & Grishchuk, E. L. Tubulin depolymerization may be an ancient biological motor. J. Cell Sci. 123, 3425–3434 (2010).

19. Gudimchuk, N. et al. Kinetochore kinesin CENP-E is a processive bi-directional tracker of dynamic microtubule tips. Nat. Cell Biol. 15, 1079–1088 (2013).

20. Gudimchuk, N. et al. Probing Mitotic CENP-E Kinesin with the Tethered Cargo Motion Assay and Laser Tweezers. Biophys. J. 114, 2640–2652 (2018).

21. Vitre, B. et al. Kinetochore-microtubule attachment throughout mitosis potentiated by the elongated stalk of the kinetochore kinesin CENP-E. Mol. Biol. Cell 25, 2272–2281 (2014).

22. Liu, D. et al. Human NUF2 interacts with centromere-associated protein E and is essential for a stable spindle microtubule-kinetochore attachment. J. Biol. Chem. 282, 21415–21424 (2007).

23. Johnston, K. et al. Vertebrate kinetochore protein architecture: protein copy number. J. Cell Biol. 189, 937–43 (2010).

24. Lawrimore, J., Bloom, K. S. & Salmon, E. D. Point centromeres contain more than a single centromere-specific Cse4 (CENP-A) nucleosome. J. Cell Biol. 195, 573–82 (2011).

25. Sundin, L. J. R., Guimaraes, G. J. & Deluca, J. G. The NDC80 complex proteins Nuf2 and Hec1 make distinct contributions to kinetochore-microtubule attachment in mitosis. Mol. Biol. Cell 22, 759–68 (2011).

26. Ciferri, C. et al. Implications for kinetochore-microtubule attachment from the structure of an engineered Ndc80 complex. Cell 133, 427–439 (2008).

27. Zaytsev, A. V. et al. Multisite phosphorylation of the NDC80 complex gradually tunes its microtubule-binding affinity. Mol. Biol. Cell 26, 1829–1844 (2015).

28. O’Brien, E. T., Salmon, E. D., Walker, R. A. & Erickson, H. P. Effects of Magnesium on the Dynamic Instability of Individual Microtubules. Biochemistry 29, 6648–6656 (1990).

29. Wood, K. W. et al. Antitumor activity of an allosteric inhibitor of centromere-associated protein-E. Proc. Natl. Acad. Sci. 107, 5839–5844 (2010).

30. Hunt, A. J. & McIntosh, J. R. The dynamic behavior of individual microtubules associated with chromosomes in vitro. Mol. Biol. Cell 9, 2857–2871 (1998).

31. Dixit, R. et al. Microtubule plus-end tracking by CLIP-170 requires EB1. Proc. Natl. Acad. Sci. 106, 492–497 (2009).

32. Bieling, P. et al. Reconstitution of a microtubule plus-end tracking system in vitro. Nature 450, 1100–1105 (2007).

33. Al-Bassam, J. et al. CLASP promotes microtubule rescue by recruiting tubulin dimers to the microtubule. Dev. Cell 19, 245–258 (2010).

34. Thorn, K. S., Ubersax, J. A. & Vale, R. D. Engineering the processive run length of the kinesin motor. J. Cell Biol. 151, 1093–1100 (2000).

35. Yardimci, H., van Duffelen, M., Mao, Y., Rosenfeld, S. S. & Selvin, P. R. The mitotic kinesin CENP-E is a processive transport motor. Proc. Natl. Acad. Sci. 105, 6016–6021 (2008).

36. Barisic, M. et al. Microtubule detyrosenation guides chromosomes during mitosis. Science 348, 799–803 (2015).

37. Block, S. M., Asbury, C. L., Shaevitz, J. W. & Lang, M. J. Probing the kinesin reaction cycle with a 2D optical force clamp. Proc. Natl. Acad. Sci. 100, 2351–2356 (2003).

38. Grishchuk, E. L. et al. Different assemblies of the DAM1 complex follow shortening microtubules by distinct mechanisms. Proc. Natl. Acad. Sci. U. S. A. 105, 6918–23 (2008).

39. Maiato, H., Gomes, A., Sousa, F. & Barisic, M. Mechanisms of Chromosome Congression during Mitosis. Biology. 6, E13 (2017).

40. Brouhard, G. J. & Rice, L. M. Microtubule dynamics: An interplay of biochemistry and mechanics. Nat. Rev. Mol. Cell Biol. 19, 451–463 (2018).

41. Monda, J. K. et al. Microtubule Tip Tracking by the Spindle and Kinetochore Protein Ska1 Requires Diverse Tubulin-Interacting Surfaces. Curr. Biol. 27, 3666–3675 (2017).

42. Hill, T. L. Theoretical problems related to the attachment of microtubules to kinetochores. Proc. Natl. Acad. Sci. 82, 4404–4408 (1985).

43. Zaytsev, A. V, Sundin, L. J. R., DeLuca, K. F., Grishchuk, E. L. & DeLuca, J. G. Accurate phosphoregulation of kinetochore-microtubule affinity requires unconstrained molecular interactions. J. Cell Biol. 206, 45–59 (2014).

44. Zaytsev, A. V, Ataullakhanov, F. I. & Grishchuk, E. L. Highly Transient Molecular Interactions Underlie the Stability of Kinetochore-Microtubule Attachment During Cell Division. Cell. Mol. Bioeng. 6, 393–405 (2013).

45. Miller, H. P. & Wilson, L. Preparation of microtubule protein and purified tubulin from bovine brain by cycles of assembly and disassembly and phosphocellulose chromatography. Methods in Cell Biology 95, 3–15 (2010).

46. Kim, Y., Heuser, J. E., Waterman, C. M. & Cleveland, D. W. CENP-E combines a slow, processive motor and a flexible coiled coil to produce an essential motile kinetochore tether. J. Cell Biol. 181, 411–419 (2008).

47. Maurer, S. P. et al. EB1 accelerates two conformational transitions important for microtubule maturation and dynamics. Curr. Biol. 24, 372–384 (2014).

48. Welburn, J. P. I. et al. The human kinetochore Ska1 complex facilitates microtubule depolymerization-coupled motility. Dev. Cell 16, 374–85 (2009).

49. Bieniossek, C., Richmond, T. J. & Berger, I. MultiBac: Multigene baculovirus-based eukaryotic protein complex production. Current Protocols in Protein Science 51, 5.20.1-5.20.26 (2008).

50. Chakraborty, M., Tarasovetc, E. V. & Grishchuk, E. L. In vitro reconstitution of lateral to end-on conversion of kinetochore–microtubule attachments. Methods in Cell Biology 144, 307–327 (2018).

51. Volkov, V. a, Zaytsev, A. V & Grishchuk, E. L. Preparation of segmented microtubules to study motions driven by the disassembling microtubule ends. J. Vis. Exp. 85, e51150 (2014).

52. Eliceiri, K., Schneider, C. A., Rasband, W. S. & Eliceiri, K. W. NIH Image to ImageJ: 25 years of image analysis. Nat. Methods 9, 671–675 (2012).

53. Andreasson, J. O. L. et al. Examining kinesin processivity within a general gating framework. Elife 4, e07403 (2015).

54. Schnitzer, M. J., Visscher, K. & Block, S. M. Force production by single kinesin motors. Nat. Cell Biol. 2, 718–723 (2000).

55. Visscher, K., Schnltzer, M. J. & Block, S. M. Single kinesin molecules studied with a molecular force clamp. Nature 400, 184–189 (1999).

56. Milic, B., Andreasson, J. O. L., Hancock, W. O. & Block, S. M. Kinesin processivity is gated by phosphate release. Proc. Natl. Acad. Sci. 111, 14136–14140 (2014).

57. Wilson-Kubalek, E. M., Cheeseman, I. M., Yoshioka, C., Desai, A. & Milligan, R. A. Orientation and structure of the Ndc80 complex on the microtubule lattice. J. Cell Biol. 182, 1055–61 (2008).

58. Walcott, S. The load dependence of rate constants. J. Chem. Phys. 128, 215101 (2008).

59. Arpa?, G., Shastry, S., Hancock, W. O. & Tüzel, E. Transport by populations of fast and slow kinesins uncovers novel family-dependent motor characteristics important for in vivo function. Biophys. J. 107, 1896–1904 (2014).

60. Lansky, Z. et al. Diffusible crosslinkers generate directed forces in microtubule networks. Cell 160, 1159–1168 (2015).

61. Jiang, L., Gao, Y., Mao, F., Liu, Z. & Lai, L. Potential of mean force for protein-protein interaction studies. Proteins Struct. Funct. Genet. 46, 190–196 (2002).

62. Lopez, B. J. & Valentine, M. T. The +TIP coordinating protein EB1 is highly dynamic and diffusive on microtubules, sensitive to GTP analog, ionic strength, and EB1 concentration. Cytoskeleton 73, 23–34 (2016).

63. Gillespie, D. T. Exact stochastic simulation of coupled chemical reactions. J. Phys. Chem. 81, 2340–2361 (1977).

