## Supplementary material for "Molecular requirements for transition from lateral to end-on microtubule binding and dynamic coupling"

**CONTENTS**

**SUPPLEMENTARY NOTE: DESCRIPTION OF THE MATHEMATICAL MODEL**

General overview of the model
Description of model parts
Description of the MT in the one-dimensional model
Description of molecular motors
Description of the diffusing MAPs
Choice of model parameters
Description of the simulation algorithm
Analysis of the simulation results
Generation of computational videos

**LEGENDS TO VIDEOS**
**TABLES S1 AND S2**
**SUPPLEMENTARY FIGURES**

### DESCRIPTION OF THE MATHEMATICAL MODEL

#### General overview of the model

A mathematical model was designed to investigate the dynamic and force-sensitive interactions between a microtubule (MT) and a multi-molecular ensemble of MT wall-binding proteins. These proteins include transporting molecular motors (kinesins) and MT-associated proteins (MAPs). The MT is modeled as a rigid rod, which is subjected to thermal motions and viscous drag. To match the geometry of our end-conversion assay, the MT-binding molecules are distributed randomly on a flat surface measuring 250 × 40 nm. The size of this patch corresponds roughly to the estimated area of the surface of a microbead, from which the molecules can reach the wall of a laterally-attached MT (Fig. 2a). A similar area of mitotic kinetochore, which has a radius of 200 nm, engages in lateral MT binding. Each molecule is firmly attached at one end to the surface of the patch, whereas its opposite end contains a MT-binding site. Molecular stalks are modeled as springs, allowing them to mediate mechanical coupling within this system. For simplicity, the maximum molecular extension length and rigidity are assumed to be the same for MAPs and motors (Table S1). The MT-interacting ends of the molecules bind and unbind stochastically from the regularly located binding sites on the MT lattice (Fig. 2b). In addition, the motors can step unidirectionally, whereas the MAP molecules can diffuse on the MT wall (Fig. 2b). The advanced feature of our model is that all of these transitions are force-sensitive, as described in detail in the following sections. Additionally, Langevin equations are used to calculate the coordinates of all molecules and MT ends, leading to a realistic and mechanically accurate description of motions and forces arising in this molecular-mechanical system.

#### Description of model parts

Because comprehensive modeling of the three-dimensional interactions involving a full MT cylinder and multiple interacting molecules would be very computationally intensive, we considered a simplified one-dimensional version of the model. In the one-dimensional representation, the 13-protofilament MT is replaced by a single-protofilament MT moving along one axis.

#### Description of the MT in the one-dimensional model

The MT contains a single protofilament 8 μm in length, which does not change during the simulation, as in our experiments with GMPCPP-containing MTs. The position of the modeled MT is represented by the coordinate of its plus-end. The molecular binding sites form a linear array along this rod with a periodicity Δ = 4 nm. To take into account the fact that in reality, the patch-immobilized molecules can interact with several protofilaments of the same MT cylinder, in our one-dimensional model we allow up to 5 molecules to simultaneously occupy a single binding site within the modeled linear array.

Three types of forces act on such MTs in our model: thermal forces of Brownian noise, viscous drag friction, and forces transmitted by MT-bound molecules. Thermal forces acting on the MT lead to random fluctuations in the MT position, calculated in each iteration as:

$$\Delta x_{thermal} = \sqrt{2\frac{k_B T}{\gamma}\,\Delta t}\; N(0,1) \qquad (1)$$

where $\Delta t$ is the time step, $\gamma$ is the viscous drag coefficient for the MT, $N(0,1)$ is a random number from a normal distribution, $k_B$ is the Boltzmann constant, and $T$ is the temperature.

Viscous drag acting on the MT is assumed to be proportional to MT velocity:

$$F_{viscous} = \gamma \, \frac{x_i - x_{i-1}}{\Delta t} \tag{2}$$

where $x_i$ and $x_{i-1}$ are consecutive MT positions separated by time $\Delta t$.

The force acting on the MT from a MT-bound molecule is calculated as molecular stiffness $k_{stiffness}$ multiplied by molecular extension.

**Description of molecular motors**

Kinesin is modeled with a single head that binds to MT sites stochastically with binding rate $k_{on}^{C(k)}$ . Subsequently, the motor translocates toward the MT plus-end with stepping rate $k_{step}^{C(k)} = V(F)/(2\,\Delta)$, where $2\,\Delta = 8$ nm corresponds to the kinesin step size, and $V(F)$ is the velocity of motor motion under force $F$. The force-dependent velocity $V(F)$ for CENP-E kinesin was taken from [20] (Table S1). Data for Kinesin-1 at 2 mM ATP is based on Figure 5 in [53]. This experimental function was fit as in [54] using the following expression:

$$V(F) = V_u\left(p_{FV} + (1 - p_{FV})e^{\frac{F\,d_{FV}}{k_B T}}\right)^{-1} \tag{3}$$

Here, $V_u$ is the unloaded motor velocity, $d_{FV}$ is the characteristic distance over which the load acts, and $p_{FV}$ and $(1 - p_{FV})$ are the fractions of biochemical and mechanical transitions in the kinesin stepping cycle, respectively.

For Kinesin-1 at for 20 µM ATP, we used the same values of $p_{FV}$ and $d_{FV}$ as for 2 mM ATP, as these parameters are not affected significantly by ATP concentration [53,55]. The unloaded motor velocity (parameter $V_u$) for Kinesin-1 at 20 µM ATP was estimated from the experimental dependency of $V_u$ on ATP concentration [37].

Additionally, the model incorporates the force-dependent unbinding of motor molecules from the MT. Data for CENP-E motor unbinding rate were taken from [20]. For CENP-E, the unbinding rate $k_{off}^{c}$ increases exponentially with force, and this increase is symmetric for assisting and opposing loads. Data for Kinesin-1 at 2 mM ATP concentration were taken from Figure 6 in [53]. Because the unbinding rate $k_{off}^{k}$ for Kinesin-1 is asymmetric with respect to the direction of applied force $F$, a linear fit was used for assisting loads ($F \geq 0$), whereas an exponential fit was used for opposing loads ($F < 0$):

$$F \geq 0:\;\; k_{off}^{k} = k_{off}^{ko} + \delta^{k_{as}} F$$

$$F < 0:\;\; k_{off}^{k} = k_{off}^{ko} e^{\frac{F\,\delta^{k_{op}}}{k_B T}} \tag{4}$$

where $k_{off}^{ko}$ is the unbinding rate for Kinesin-1 without any load, $\delta^{k_{as}}$ is the force sensitivity parameter for an assisting load, and $\delta^{k_{op}}$ is the characteristic distance parameter for an opposing load.

To calculate the force-unbinding function for Kinesin-1 at 20 µM ATP, we took into account the fact that the run length of Kinesin-1 is independent of ATP over a wide range of concentrations (from 2 µM to 2 mM [56]). Therefore, at low ATP concentration:

$$k_{off}^{c(k)}\Big|_{low\ ATP} = k_{off}^{c(k)} \times \frac{V(F)|_{low\ ATP}}{V(F)} \qquad (5)$$

where the *low ATP* subscript indicates the unbinding rate and force-velocity dependency at 20 µM ATP.

Finally, for all motors and conditions, the unbinding rate of the motor from the terminal binding site at the MT plus-end is assumed to be the same as its unbinding rate from the MT wall. Therefore, after the motor molecule reaches the MT plus-end, it stays bound to the end on average for the same time as it would have if it had continued walking on the MT. The unbinding from the end takes place according to the same force-dependent function as elsewhere along the MT.

**Description of the diffusing MAPs**

The distance between the MT-binding sites for the Ndc80 complex is Δ = 4 nm [57]. For simplicity, the same step size is used in the model for all other MAPs. To describe force-dependent transitions during a MAP's diffusion along the MT rod, we assume Bell's relationship [58]:

$$k^{+(-)} = D_o\, \Delta^{-2}\, e^{(-)\frac{F\,\delta}{k_B T}} \qquad (6)$$

Here, $k^+$ and $k^-$ are the MAP's stepping rates towards the MT plus or minus end, respectively; $D_o$ is the diffusion coefficient along the MT for the freely diffusing MAP in the absence of external force, as measured using single-molecule TIRF-visualization *in vitro*; $F$ is the force acting on this MAP molecule through the MT due to the MT's thermal motion, kinesin walking, and hindrance from other MT-bound MAP molecules; $\delta$ is the force-sensitivity parameter; and Δ = 4 nm is the MAP's step size.

An analogous force-dependency function was assumed for the MAP's unbinding rate from the MT wall $k_{off}^{M}$:

$$k_{off}^{M} = k_{off}^{Mo}\, e^{\frac{|F|\delta}{k_B T}} \qquad (7)$$

Here, $k_{off}^{Mo}$ is the MT unbinding rate for the freely diffusing MAP, and $|F|$ is the absolute value of the force acting on the MAP through the bound MT.

If a MAP reaches the last binding site at any MT end, we assume that it can detach with the same force-dependent rate as elsewhere on the MT wall ($k_{off}^{M}$). However, unlike the motor molecule, if the MAP does not detach immediately from the terminal binding site, it does not remain motionless at that site, as the motor does. Instead, the MAP continues to diffuse along the MT (Fig. S4d). From the terminal MT-binding site, the MAP can make a diffusional step towards the adjacent site with stepping rate $k^-$ if the MAP is at the MT plus-end and $k^+$if the MAP is at the minus-end. However, analogous diffusional steps in the wrong direction will obviously lead to a detachment. Thus, the total unbinding rate for a MAP from the terminal binding site at any MT end is:

$$k_{off\_end}^{M} = k_{off}^{M} + \alpha\, k^{+(-)} \qquad (8)$$

Here, $\alpha$ is a coefficient corresponding to the probability that the MAP will take a diffusional step away from the MT, leading to detachment.

Simulations for different MAPs were performed using the same dependencies, simulation algorithm (below), and model parameters except for two single-molecule characteristics: the rate of MT wall diffusion and residence time.

**Choice of model parameters**

All model parameters and corresponding values are listed in Tables S1 and S2. An additional description is provided below.

*Molecular stiffness* $k_{stiffness}$ was assumed to depend on the extension length of the molecule. When molecular length exceeded 30 nm (the approximate contour length of the Ndc80 protein complex used in our *in vitro* work), high molecular stiffness of 2,000 pN/µm was assumed to prevent significant extension beyond this length under force. For shorter extensions, stiffness was assumed to be 10-fold lower, 200 pN/µm, similar to the value used in some other models featuring molecular-MT coupling [59,60]. In our one-dimensional models, this relatively low stiffness at lengths that are smaller than the resting length enables changes in the size of the linear projection of the MT-binding molecule onto the MT axis. In a real situation, the molecule can be located some distance away from the MT (Fig. 2a), so its linear projection can be much smaller than the contour length even in the absence of force. In our model, the change in the size of a linear projection takes this geometrical aspect into account, rather than corresponding to the true extension/compression under force.

*Force-sensitivity parameter $\delta$ for MAPs* was set to 0.2 nm, within the typical range of characteristic distances for protein–protein interactions[61].

*Number of molecular motors and MAPs interacting with the MT wall.* To estimate the number of MAPs bound to the MT during our end-conversion assays, we took advantage of the relationship between the number and diffusion rate of the individual MAPs and the resultant diffusion rate of the bound MT: when more molecules are bound, MT diffusion is slower. To define this relationship quantitatively, we carried out simulations using our model with different numbers of patch-bound MAPs (Fig. S3c). In these simulations, we used $D_o$ = 0.089 µm$^2$/s , corresponding to the diffusion coefficient of one Ndc80 molecule, which we determined *in vitro* for Ndc80-Broccoli on GMPCPP MTs (Table S2). The unbinding rate for the MAPs in these simulations was assumed to be 0 to keep constant the number of the MT-bound molecules, which is therefore equal to the total number of MAPs on the patch. The resultant MT diffusion coefficients were plotted, revealing that MT diffusion coefficient decreases hyperbolically as the number of MAPs increases (Fig. S3d).

We also derived the analytical dependency for this relationship. If all molecules within a molecular patch step independently, the stepping rate of the molecular patch is $Nk^o$, where $N$ is the number of MAPs in the molecular patch, and $k^o = D_o\Delta^{-2}$ is the stepping rate of a single molecule. Displacement of the molecular patch during one step is $\Omega = \Delta/N$, where $\Delta$ is the step size for a single MAP. Therefore, the diffusion coefficient $D$ of the molecular patch is given by:

$$D = \Omega^2 k = \frac{\Delta^2}{N^2} N k^o = \frac{1}{N}\Delta^2 k^o = \frac{1}{N} D_o \quad (9)$$

This hyperbolic function is plotted on Fig. S3d (red curve) and shows a good match to numerical simulation results (black dots). These theoretical results were then compared with the MT diffusion coefficient measured *in vitro* using coverslip-immobilized beads coated with the Ndc80 protein: 5.6 · $10^{-3}$ $\mu m^2/s$. This measured value implies that 11–13 Ndc80 complexes were bound to the MT during its diffusion (Fig. S3d, grey lines). Importantly, in our simulations of MT end-retention, the number of MT-bound molecules is less than the total number of Ndc80 molecules immobilized on the molecular patch because their unbinding rate is not 0, as we assumed in simplified calculations for Fig. S3c,d. End-retention simulations were carried out using total number of Ndc80 molecules $N_{MAPs}$ = 15 (Table S1), because this value generates the number of MT-bound Ndc80 molecules in the 11-13 range. The same number $N_{MAPs}$ was assumed when modeling other MAPs, as the brightness of beads coated with different MAPs was approximately similar in our *in vitro* end-conversions assays (Fig. S6a). Bead coating with motors was ~3-fold brighter than with the MAPs (Fig. S6a). Therefore, the number of motors in the patch in our model was $N_{motors} = 45$.

*Parameters of MAP diffusion on the MT wall* (Table S2). MT-wall diffusion coefficients and residence times for Ndc80, Ska1, and CLASP2 were measured using single-molecule TIRF microscopy *in vitro* under the same experimental conditions as in end-conversion assay (Fig. S3e-f). The experimental conditions in the published studies of the diffusion of EB1 [62] and CENP-E Tail [19] were also similar.

*Parameter $\alpha$ for the detachment of MAPs from the terminal MT-binding site.* The value of this parameter for Ndc80 and other MAPs is not known. When $\alpha$ < 1, the probability that the MAP will take a step leading to detachment is smaller than the probability of the analogous step between the adjacent MT-binding sites on the MT wall during diffusion, leading to an overall longer MT association. To investigate this effect in more details we calculated the end-retention survival probability plots for different values of α using simulation parameters corresponding to Ndc80 and CENP-E Tail (Table S2). We then calculated the difference between the predicted percent of survived MT ends at 30 min from the start of simulations for these two MAPs. With increasing α, this difference increased and came to a plateau at $\alpha = 10^{-2}$, implying that model predictions for end-retention are no longer sensitive to this parameter (Fig. S4e). Accordingly, this value was used for simulations with all remaining MAPs. Relatively good agreement between the resultant predictions and experimental measurements for these MAPs suggest that the value of $\alpha$ is indeed similar for these proteins. The mismatch for Ska1 complex could be explained by a different value of $\alpha$ for this protein or more trivial factors, e.g., partial inaccessibility of the Ska1 microtubule-binding site after it is conjugated to the bead surface. This would lead to a shorter end-retention time in experiment than in the model, which does not include such complexities. Future work using single-molecule visualization at MT ends is needed to determine the experimental values of $\alpha$ for different MAPs.

**Description of the simulation algorithm**

At the start of each calculation, the positions of all MAPs and kinesin motors along the patch-representing linear segment were chosen randomly. The initial MT configuration was centered relative to this segment, and calculations started with all molecules not bound to the MT. Subsequent iterations were carried out with the time step $\Delta t$ for total simulation time $t_{total}$. For each stochastic event *E* occurring with rate *k*, the probability $\Psi_E$ of this event to occur at each time step was calculated as in [63]:

$$\Psi_E = 1 - e^{-\Delta t\, k} \tag{10}$$

Next, a random number *p* from the range [0, 1] with constant probability density was generated. If *p* was smaller than $\Psi_E$, the corresponding stochastic event *E* was assumed to be accomplished.

Four types of stochastic events were calculated at each step:

(1) Binding of MAPs and motors to sites on the MT. For each unbound molecule, the linear distance between its surface-attached end and the nearest binding site on the MT was calculated. If the distance to the nearest MT site was ≤ 2 nm, the MAP (or motor) molecule was assumed to bind to this site with rate $k^{M}_{on}$ (or $k^{k}_{on}$).

(2) Unbinding of molecular motors and MAPs from the MT. For each MT-bound molecule, the applied force $F$ was calculated as molecular stiffness $k_{stiffness}$ multiplied by the molecular extension length, calculated as the linear distance between the surface-bound and MT-bound ends of the molecule (corresponding to the length projection in three-dimensional space). Each MT-bound molecule detached according to force-dependent unbinding rates, as described in the sections for MAPs and kinesins.

(3) MT-dependent translocation of molecules. The stepping of kinesin motors and diffusion of MAPs was calculated according to the force-dependent transition rates described in the corresponding sections.

(4) Motion of the MT. The coordinate of the new MT plus-end position $x_i^{MT}$ was calculated using the Langevin equation:

$$x_i^{MT} = x_{i-1}^{MT} + \frac{\Delta t}{\gamma} F_{total} + \sqrt{2\frac{k_B T}{\gamma}}\, \Delta t\, N(0,1) \qquad (11)$$

where $x_{i-1}^{MT}$ is the MT plus-end position at the previous time step, $\Delta t$ is the time step, $\gamma$ is the viscous drag friction coefficient for the MT, $N(0,1)$ is a random number from a normal distribution, $k_B$ is the Boltzmann constant, $T$ is the temperature, and $F_{total}$ is the total force applied to the MT from all MT-bound MAPs and motors.

$F_{total}$ was calculated using the following expression:

$$F_{total} = \sum_{m=1}^{N_{boundMAPs}} \Delta l_m k_{stiffness} + \sum_{n=1}^{N_{boundKinesins}} \Delta l_n k_{stiffness}^{Kinesin} \qquad (12)$$

where $\Delta l_m$ corresponds to the extensions of each MAP bound to the MT, and $\Delta l_n$ corresponds to the extensions of each kinesin bound to MT; and $N_{boundMAPs}$ and $N_{boundKinesins}$ are the total numbers of MT-bound MAPs and motors at this time step, respectively.

After these stochastic events were calculated and MT position was updated, the positions of all binding sites on the MT relative to the surface-bound molecules were also updated, yielding the updated molecular extension lengths and forces acting on each molecule. The simulation time was increased by $\Delta t$ and compared with $t_{total}$: if the simulation time was less than $t_{total}$, the algorithm was repeated from step (1); if the simulation time was greater than $t_{total}$, the simulation was stopped.

**Analysis of the simulation results**

To obtain the end-retention survival probability plot from model simulations, we carried out simulations for $t_{total}$ = 30 min, corresponding to the duration of experimental observations. Survival time was defined as the first-time step from the start of simulations for which the MT had zero bound MAPs and motors. The end-retention survival probability curve was then plotted from the fraction of MTs that had at least one bound molecule (either a motor or a MAP) at each time step, based on 32 independent simulations for each pair of motors and MAPs.

**Generation of computational videos**

To generate three-dimensional computational videos, calculations were carried out using a one-dimensional model, as described in the simulation algorithm. Then, image frames were generated using a custom-designed program written in Mathematica to represent three-dimensional interactions. Briefly, a single protofilament was complemented with tubulin-representing spheres to form a full cylinder 0.9 µm long, and then positioned 0.02 µm above the patch surface. Positions of the surface-bound ends of each molecule along the MT axis were as in the original one-dimensional simulation, whereas their positions in the MT-perpendicular direction were chosen randomly within a 45-nm distance from the protofilament, mimicking a two-dimensional patch with attached molecules. Translocation of each molecule along the MT was assumed to take place along the nearest MT protofilament, and the coordinates of the MT-bound ends of all molecules at each time point were taken from the corresponding one-dimensional simulation. To improve visual discrimination of the MT-bound and -unbound molecules, the latter are shown in the videos in random orientations. Furthermore, to avoid cluttering, the number of molecules in these simulations was reduced relative to our standard simulations: $N_{MAPs}$ = 5, $N_{motor}$ = 5. Other parameters used in video simulations: $k_{on}$ = 3.6 $s^{-1}$, $k_{on}$ = 120 $s^{-1}$, MAP diffusion coefficient $D_o$ = 0.0038 $\mu m^2/s$.

### LEGENDS TO VIDEOS

**Video 1. MT gliding on beads coated with CENP-E motors.**
In this and all other experimental videos, the GMPCPP-stabilized MTs are labeled with Hilyte-675, so they appear in red; green indicates coverslip-immobilized 1-µm beads coated with GFP-tagged proteins. Scale bar, 3 µm. In this video, beads are coated with the motor domains of CENP-E. MTs are initially attached laterally, but when ATP is added they glide on the beads and detach quickly. Because CENP-E is a plus-end–directed motor, the MT plus-end is the last contact point between the gliding MT and the bead. Video plays 65 times faster than actual speed.

**Video 2. MT wall-to-end transition by CENP-E kinesin and Ndc80 complex.**
MTs glide visibly more slowly on beads coated with a mixture of CENP-E motors and Ndc80 Broccoli than on beads coated with CENP-E alone (compare to Video 1). After the MT plus-end reaches the bead, it is retained rather than dissociating, often persisting for longer than the total imaging time. Video plays 65 times faster than actual speed.

**Video 3. Modeling of MT gliding over a molecular patch with CENP-E motors.**
In this and all other computational videos, MT is depicted in green, motors in blue, and Ndc80 molecules in red; for ease of viewing, only five molecules of each type are included. The video frames playing at 30 fps show results for one out of every $10^4$-time steps of the corresponding simulation, yielding motions 3.3-fold slower than those in the model. In this video, the patch-immobilized motors bind randomly and step toward the MT plus-end, propelling the MT minus-end forward. When the MT plus-end arrives at the molecular patch, it pauses briefly, awaiting the dissociation of the last motor.

**Video 4. Modeling of MT diffusion over a molecular patch with Ndc80 molecules.**

Patch-immobilized Ndc80 molecules randomly bind, diffuse on the MT wall, and then unbind, collectively driving the MT diffusion. MT ends are sometimes seen in the vicinity of the patch, but on average the MT is centered because all interactions are thermally driven, and the motions are not biased.

**Video 5. Modeling of MT wall-to-end transition by CENP-E and Ndc80.**
This calculation combines CENP-E motors and Ndc80 molecules, which interact with the MT in essentially the same manner as in Videos 3 and 4. Their force-dependent ensemble behavior leads to durable MT end retention, even though the model assumes no special interactions between these molecules and MT tips.

**Video 6. Dynamic coupling between MT plus-ends and the bead-immobilized CENP-E and Ndc80 proteins.**
The video starts after unlabeled soluble tubulin and GTP are added to the chamber, in which brightly labeled GMPCPP-stabilized MTs were transported and underwent end attachment to beads coated with CENP-E and Ndc80. Two labeled MT segments are seen slowly moving away from the bead and quickly retracting back, implying that tubulin is incorporated and lost from the bead-bound MT plus-ends. Video plays 30 times faster than actual speed. Scale bar, 3 µm.

**Video 7. Dynamic coupling between MT plus-ends and bead-immobilized CENP-E and Ndc80 proteins in high $MgCl_2$.**
This video is analogous to Video 6, but the experiment was performed in a buffer containing 7 mM instead of 2 mM $MgCl_2$. The rate at which the bright MT segment moves away from the bead is elevated, consistent with a higher rate of tubulin incorporation at the bead-coupled MT plus-end. Video plays 30 times faster than actual speed. Scale bar, 3 µm.

**Video 8. MT wall-to-end transition by CENP-E kinesin and the Ska1 complex.**
A GMPCPP-stabilized MT glides on a bead coated with a mixture of CENP-E motors and Ska1 proteins, undergoing wall-to-end transition. However, the MT plus-end detaches several minutes after establishing end attachment, indicating a problem with maintenance. Video plays 65 times faster than actual speed. Scale bar, 3 µm.

**Video 9. MT wall-to-end transition by CENP-E kinesin and EB1 protein, reverse end-to-wall transition, and lateral MT diffusion.**
GMPCPP-stabilized MTs glide on a bead coated with a mixture of CENP-E motors and Ska1 proteins, undergoing wall-to-end transition. After a few minutes, one of these MTs (labeled MT1) loses end attachment and undergoes lateral diffusion. CENP-E then returns the MT plus-end to the bead, and end coupling is re-established. Video plays 65 times faster than actual speed. Scale bar, 3 µm.

**Video 10. MT wall-to-end transition by Kinesin-1 and Ndc80 proteins.**
GMPCPP-stabilized MTs bind to beads coated with a mixture of Kinesin-1 motors and Ndc80 proteins, glide rapidly, and detach almost immediately, indicating that they are unable to maintain end attachment. Video plays 65 times faster than actual speed. Scale bar, 3 µm.

**TABLE S1. PARAMETERS USED IN THE SIMULATIONS**

| **General model parameters** | | | |
|---|---|---|---|
| **Symbols** | **Description** | **Value** | **Comments** |
| $\Delta t$ | simulation time step | 4 µs | chosen for convergence of simulation algorithm |
| $t_{total}$ | total simulation time | 30 min | as in experiments *in vitro* |
| $L_{MT}$ | length of microtubule | 8 µm | typical MT length in experiments *in vitro* |
| $\gamma$ | viscous drag coefficient per microtubule length | 0.014 pN·s·µm$^{-2}$ | calculated based on dynamic viscosity (0.002 Pa·s) and diameter of MT (25 nm) |
| $k_{stiffness}$ | stiffness of MAPs and motors | extension ≤ 30 nm: 200 pN/µm<br>extension > 30 nm: 2,000 pN/µm | see section "Choice of model parameters" in the Supplementary Note |
| $k_B T$ | energy scale factor | 4.11 pN·nm | |
| **Parameters describing MAPs** | | | |
| **Symbols** | **Description** | **Value** | **Comments** |
| $D_o$ | diffusion coefficient for single MAP molecule on the MT | varied to model different MAPs | measured *in vitro* data (see Table S2) |
| $\tau$ | MAP residence time on the MT during diffusion | varied to model different MAPs | measured *in vitro* (see Table S2) |
| $\Delta$ | MAP step size during diffusion | 4 nm | assumed to be the same for all MAPs |
| $\delta$ | force-sensitivity parameter | 0.2 nm | see section "Choice of model parameters" in the Supplementary Note |
| $k_{on}^{M}$ | rate of MAP binding to the MT | 20 s$^{-1}$ | see section "Choice of model parameters" in the Supplementary Note |

| | | | | | |
|---|---|---|---|---|---|
| $\alpha$ | probability that the MAP will take a diffusional step away from the MT at terminal binding site | $10^{-2}$ | | | see section "Choice of model parameters" in the Supplementary Note |
| $N_{MAPs}$ | number of MAPs in the molecular patch | 15 | | | see section "Choice of model parameters" in the Supplementary Note |
| $k_{off}^{Mo}$ | MAP unbinding rate from the MT wall without external force | | | | $1 / \tau$ |
| $k^{+(-)}$ | MAP stepping rate towards the MT plus (minus) end | | | | see eq. (6) in the Supplementary Note |
| $k_{off}^{M}$ | MAP unbinding rate from the MT wall under external force | | | | see eq. (7) in the Supplementary Note |
| $k_{off\_end}^{M}$ | MAP unbinding rate from the MT terminal binding site | | | | see eq. (8) in the Supplementary Note |
| **Parameters describing motors** | | | | | |
| **Symbols** | **Description** | **Value** | | | **Comments** |
| | | CENP-E | Kinesin-1 (2 mM ATP) | Kinesin-1 (20 μM ATP) | |
| $V_u$ | velocity of unloaded motor | 0.26 μm/s | 0.62 μm/s | 0.1 μm/s | see "Description of molecular motors" section in the Supplementary Note |
| $d_{FV}$ | force-sensitivity parameter | 2.8 nm | 5.3 nm | 5.3 nm | |
| $p_{FV}$ | fraction of biochemical transitions in kinesin stepping cycle | 0.58 | 0.99 | 0.99 | |
| $k_{on}^{C(k)}$ | rate of motor binding to the MT | $0.4\ s^{-1}$ | | | see section "Choice of model parameters" in the Supplementary Note |
| $k_{step}^{C(k)}$ | motor stepping rate toward the MT plus-end | | | | $V(F)/(2\,\Delta)$ |
| $k_{off}^{Co}$ | unbinding rate for CENP-E without external force | $0.12\ s^{-1}$ | | | Gudimchuk et al., 2018 |

| | | | |
|---|---|---|---|
| $\delta^{C}$ | force-sensitivity parameter for CENP-E unbinding | 2 nm | |
| $k_{off}^{ko}$ | unbinding rate for Kinesin-1 without external force | 1.1 $s^{-1}$ for 2 mM ATP<br>0.11 $s^{-1}$ for 20 µM ATP | |
| $\delta^{k_{op}}$ | force-sensitivity parameter for Kinesin-1 unbinding under opposing load | 0.6 nm | see "Description of molecular motors" section in the Supplementary Note |
| $\delta^{k_{as}}$ | force-sensitivity parameter for Kinesin-1 unbinding under assisting load | 1.5 $pN^{-1}\cdot s^{-1}$ for 2 mM ATP<br>0.15 $pN^{-1}\cdot s^{-1}$ for 20 µM ATP | |
| $N_{motors}$ | number of motors in the molecular patch | 45 | see section "Choice of model parameters" in the Supplementary Note |

### TABLE S2. MOLECULAR PARAMETERS OF MAP-MT INTERACTIONS MEASURED USING SINGLE-MOLECULE TIRF MICROSCOPY.

N – number of independent experiments; n – number of single molecules analyzed. For experiments with N ≥ 3, errors are SEM; for N = 2, errors are 95% confidence intervals.

| MAP | Ndc80 | Ska1 | CLASP2 | CENP-E Tail† | EB1¶ |
|---|---|---|---|---|---|
| Diffusion coefficient (*D*), $\mu m^2/s$ | 0.089 ± 0.003<br>(N=2, n=1037) | 0.23 ± 0.01<br>(N=3, n=2018) | 0.33 ± 0.02<br>(N=4, n=558) | 1.6 | 0.31 ± 0.01 |
| Residence time ($\tau$), s | 0.43 ± 0.01<br>(N=2, n=551) | 0.61 ± 0.02<br>(N=2, n=974) | 0.65 ± 0.01<br>(N=2, n=518) | 0.47 ± 0.03 | 0.08 ± 0.01 |

† data from [63]

¶ data from [39]

### SUPPLEMENTARY FIGURES

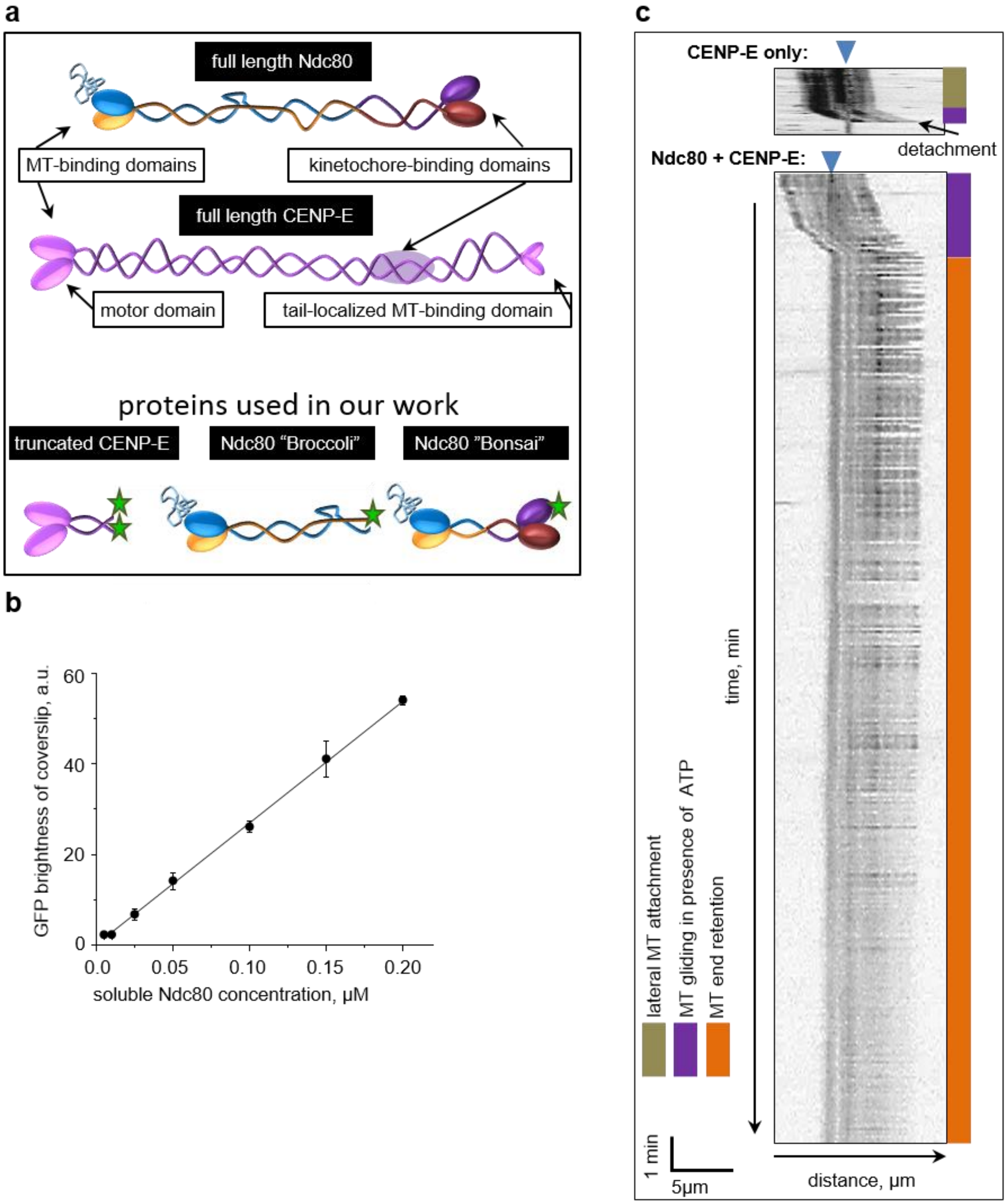


**Figure S1. MT gliding on coverslips and beads, and example kymographs**

(a) Schematic of the full-length Ndc80 and CENP-E proteins, and the truncated protein constructs used in this study.

(b) Quantification of coverslip brightness in experiments that used a mixture of proteins: Myc-tagged CENP-E with no fluorescent label and GFP-labeled Ndc80 Bonsai. The latter protein was added to the microscopy chamber at the indicated concentrations; the unbound proteins were removed; and the GFP fluorescence brightness of the coverslip coating was measured. Data points and error bars indicate means ± SEM of $N = 3$ independent experiments.

(c) Representative kymographs of motions of bead-bound stabilized MTs in the presence of ATP. Arrowheads indicate positions of coverslip-immobilized beads. Colored bars provide interpretations for the kymographs (see labels).

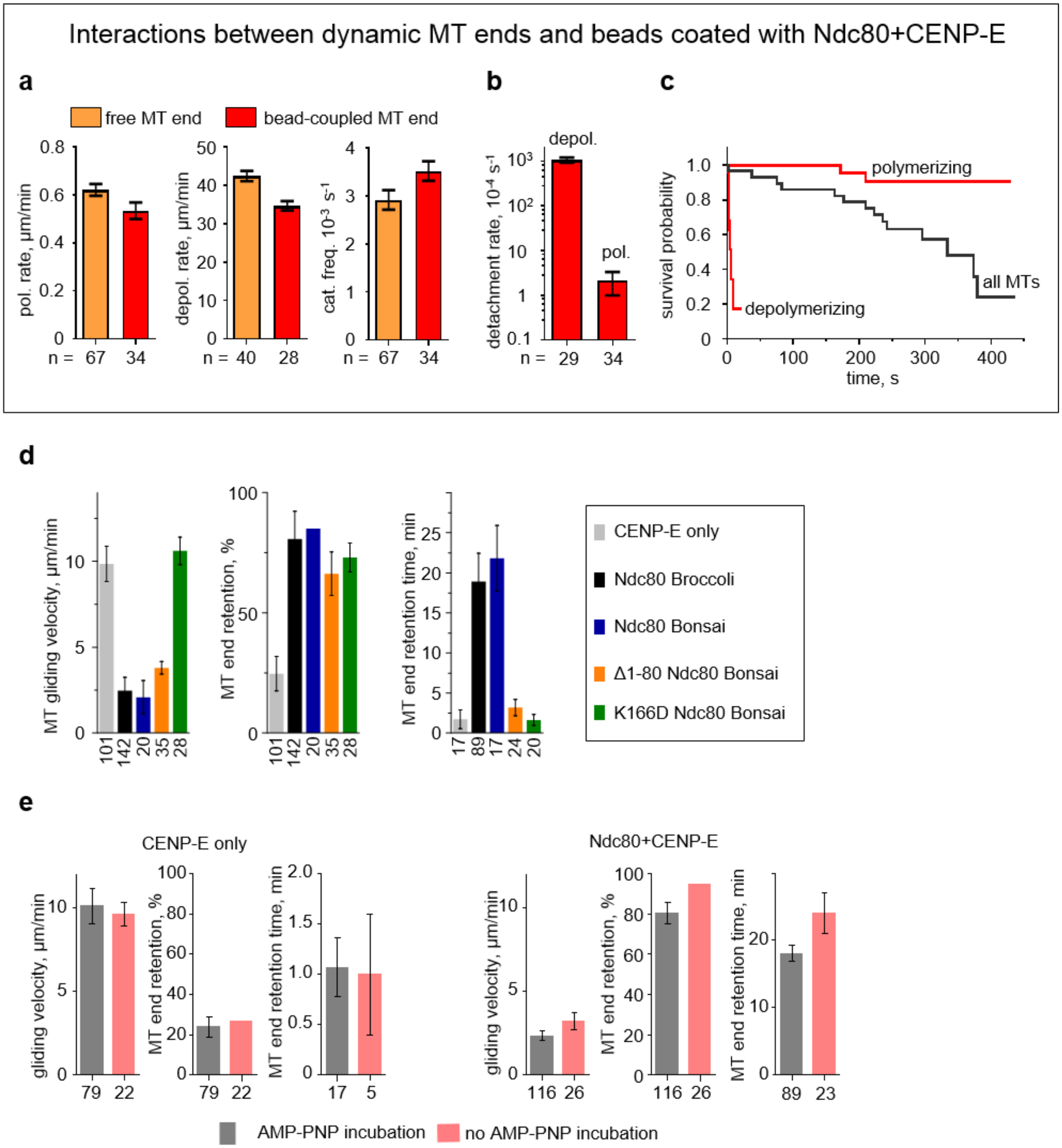


**Figure S2. Behavior of coupled MT ends in assays with floating and coverslip-immobilized beads.** Panels a–c show results for MT end coupling in experiments using freely floating Ndc80+CENP-E beads (in the 1:4 ratio) and MTs grown from coverslip-immobilized seeds. In these experiments free floating beads randomly attached to MTs, transported to the MT end by CENP-E motor and travelled at the end of dynamic MTs. Panels d and e show results for the end-conversion assays using coverslip-immobilized beads.

(a) Comparison of the dynamics of MT ends that are coupled to the beads or not coupled (i.e., free MT ends). Columns and error bars indicate means ± SEM of total number of examined MT ends (indicated under each column) for 2 independent experiments. To estimate SEM for catastrophe frequency, bootstrapping was performed using Excel: data from all MTs were pulled together, and 10 random points were drawn with replacement 10 times. These graphs show that presence of the beads does not affect significantly the rates of MT polymerization and depolymerization, as well as the catastrophe frequency.

(b) Frequency of bead detachment from the ends of dynamic MTs was calculated as inverse of the ratio of total time beads were bound at the MT ends to the number of detachment events, calculated for the total number of examined MT ends (indicated under each column). SEM was estimated as for catastrophe frequency in panel a. This graph shows that bead coupling is more stable at polymerizing MT ends than on depolymerizing ends.

(c) Survival plot of attachment time for beads coupled to the MT ends. Graphs are based on the same data set as in panel (b). Consistent with data in panel (b), bead coupling is more durable at the polymerizing MT ends.

(d) Results for end-conversion assays using a mixture of CENP-E motors and mutant Ndc80 proteins. Data for “Ndc80 Broccoli” and “CENP-E only” are the same as in Fig. 1g. Each condition was examined in at least two independent trials, yielding *n* MTs, as indicated below each column. Data are means ± SEM.

(e) Results from experiments in which stabilized MTs bound to coverslip-immobilized beads in the presence of AMP-PNP, which was subsequently replaced with ATP to initiate gliding (grey columns show means ± SEM, $N \geq 3$; total number of examined MTs is shown under each column). Some MTs attached to beads via random collisions in the presence of ATP and became immediately transported (pink columns show mean ± SEM for the number of MTs indicated under each column). Because no significant differences were observed between these different assay conditions, these data sets were combined.

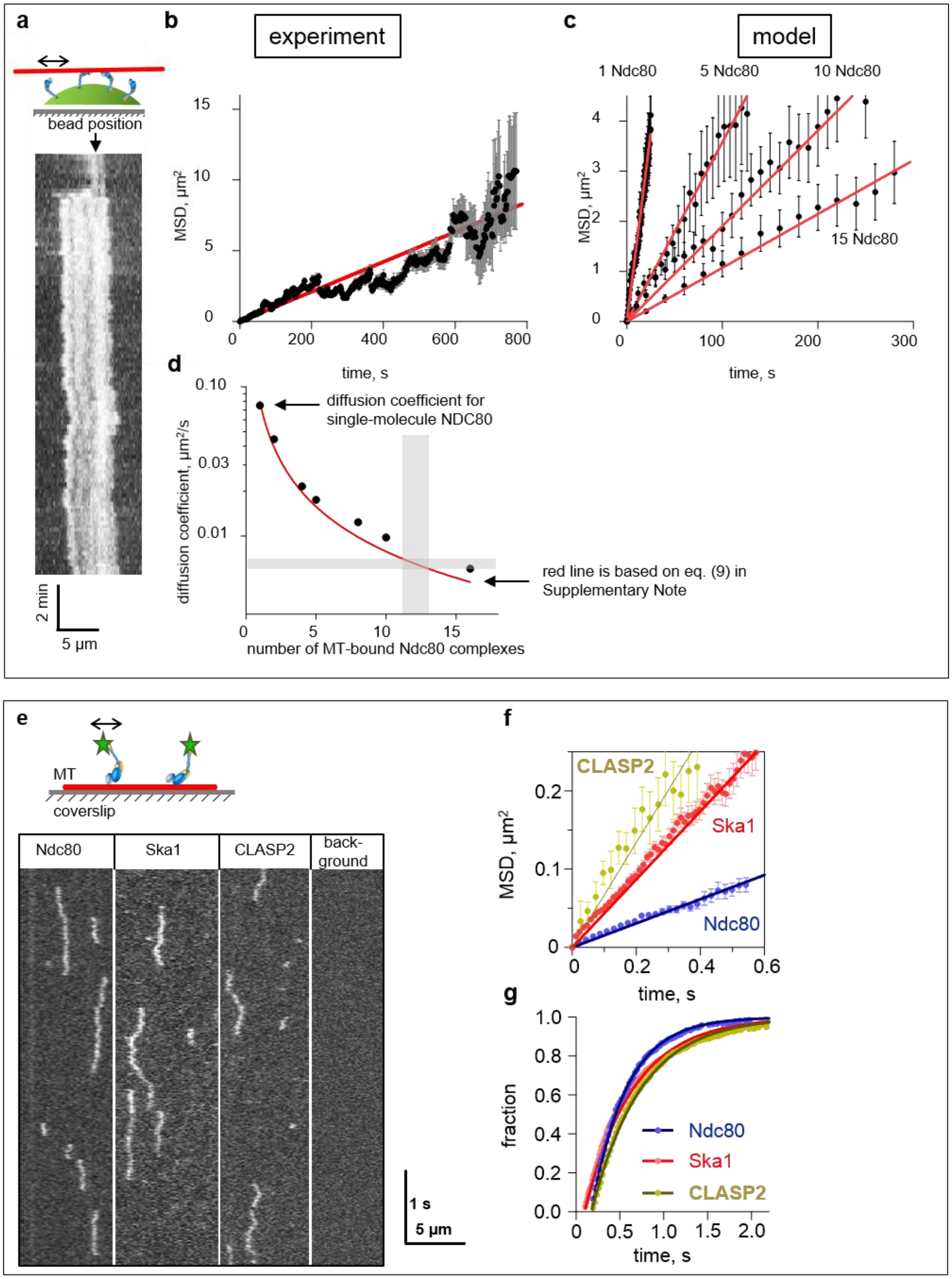


**Figure S3. Analysis of the diffusion of MTs and single molecules.**
(a) Schematic of an experiment in which stabilized MTs diffused on the surface of Ndc80-coated beads. Representative kymograph shows one diffusing MT. The position of the coverslip-immobilized bead is marked with a black arrow.

(b) Mean-square displacement (MSD) of diffusing MTs vs. time (based on tracings of *n* = 20 MTs observed in *N* = 2 independent experiments). Red line is the linear fit. Error bars are SEM.
(c) Theoretical predictions of MTs diffusing on molecular patches with different numbers of MT-bound Ndc80 molecules. Error bars are SEM.
(d) Theoretical predictions of the relationship between diffusion coefficient and the number of MT-bound Ndc80 molecules. Black dots are data from numerical simulations, and the red line is the prediction based on analytical considerations. Shaded horizontal bar corresponds to the experimentally measured MT diffusion coefficient. Shaded vertical bar corresponds to the estimated range of the number of MT-bound Ndc80 molecules.
(e) Representative kymographs depicting diffusion of single molecules of indicated MAPs, visualized via TIRF microscopy. Background image was obtained in an area with no MTs.
(f) MSD of single-molecule diffusion. Error bars are SEM.
(g) Cumulative distributions of residence times during diffusional events for the indicated MAPs.

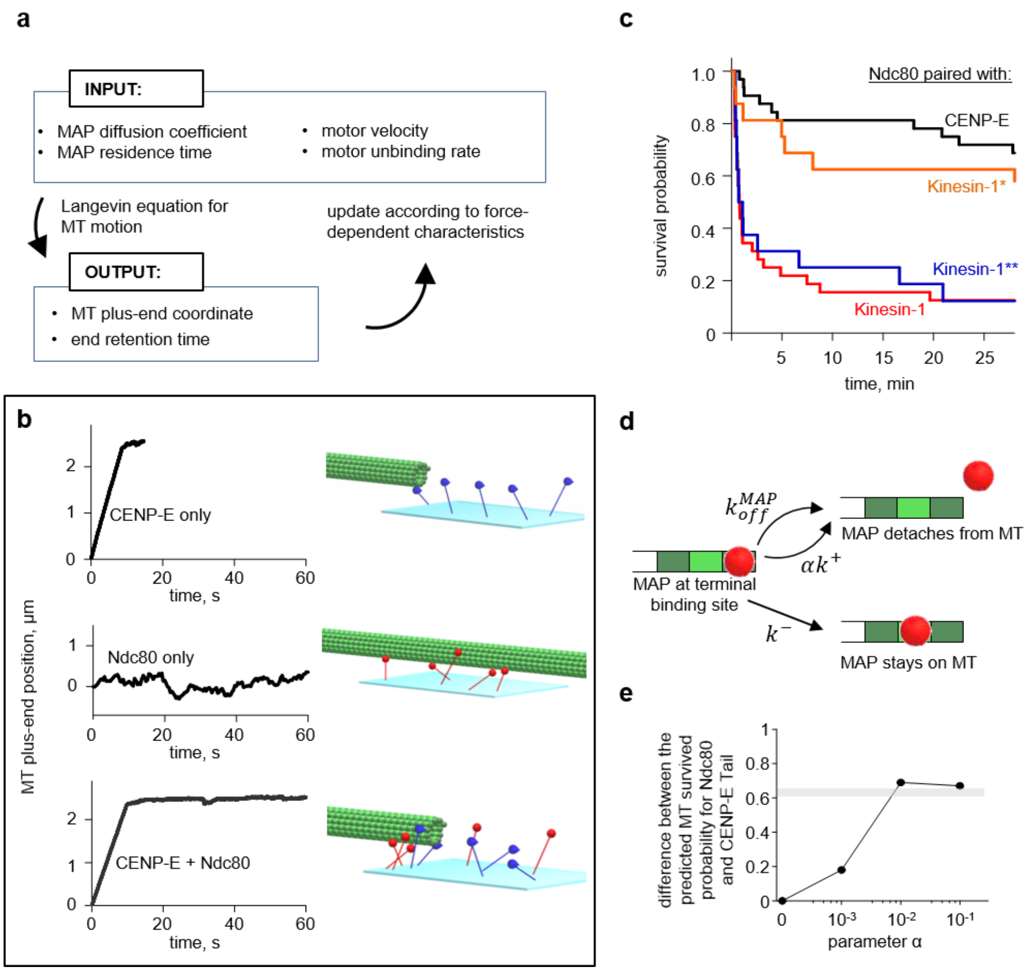


**Figure S4. Mathematical modeling of MT end conversion.**

(a) Schematic of computational flow in the model. Input parameters and dependencies were used to calculate the coordinates of MT plus-ends. Forces acting on each MAP and motor molecules were then calculated to determine the probabilities of molecular stepping (diffusional or directional) and unbinding from the MT.

(b) Representative trajectories for MT plus-ends (left) calculated for molecular patches with different molecular compositions (right).

(c) Survival probability plots calculated for molecular patches containing Ndc80 molecules and motor molecules with different force-dependent characteristics. Predictions for Ndc80+CENP-E and Ndc80+Kinesin-1 are the same as in Fig. 5e. Prediction for Kinesin-1* was calculated using force-dependent unbinding rate of CENP-E and the force-velocity function of Kinesin-1 motor. Prediction for Kinesin-1** was calculated using the force-dependent unbinding rate of Kinesin-1 and the force- velocity function of CENP-E.

(d) Schematic illustration of the behavior of a MAP molecule at the terminal binding site of the MT. The molecule can step away from the MT end, as during wall diffusion, or can detach via two independent pathways (see model description for details).

(e) MT end-retention outcome as a function of the parameter $\alpha$ (see "Choice of model parameters" in the Supplementary Note for details).

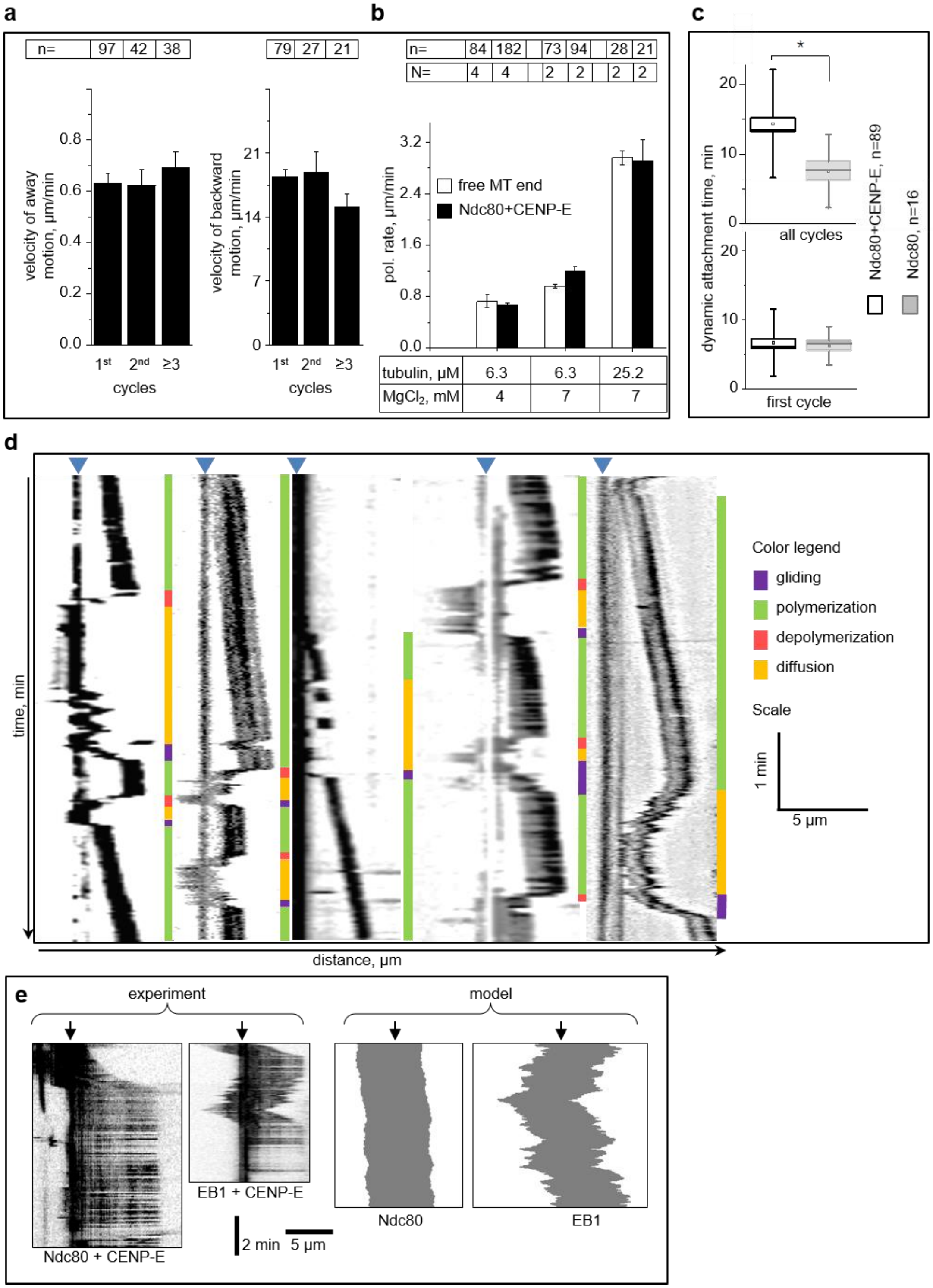


**Figure S5. Dynamics of coupled MT ends.**

(a) Results from an end-conversion assay using Ndc80+CENP-E beads and soluble tubulin. The graph shows the velocity with which labeled MT fragment moved away and toward the bead. Columns are medians ± SEM for data from *N* = 4 independent experiments with *n* observed MTs, as indicated above each column. Velocities were similar during repeated MT dynamics cycles, consistent with the idea that tubulin dynamics are the driving force.

(b) Rate of polymerization of bead-free MT plus-ends and the velocity with which labeled MT fragments moved away from beads coated with Ndc80 and CENP-E. Columns are means ± SEM for N=4 independent experiments and means ± SD (determined by bootstrap statistical analysis) for N=2 experiments. Concentrations of soluble tubulin and $MgCl_2$ are indicated below each column.

(c) Duration of dynamic attachment for the first cycle or all cycles of MT dynamics for beads coated with Ndc80, with or without CENP-E. Box is median ± SEM and whiskers are SDs; data are combined from N=4 (Ndc80+CENP-E) or N=6 (Ndc80) independent trials. Significance of differences between two data sets was evaluated using the Mann–Whitney test; * = $p < 0.05$. The duration of the first cycle is same for these bead coatings, but the total duration was significantly reduced in the absence of CENP-E.

(d) Example kymographs of dynamic MT ends coupled to beads coated with EB1+CENP-E. Colored bars provide interpretations for the kymographs (see color legend).

(e) Analysis of MT diffusion following the loss of MT tip attachment (i.e. end-to-wall transition) to the coverslip-immobilized bead. Experimental kymographs show MTs gliding on the beads coated with CENP-E and either Ndc80 or EB1. The latter MT subsequently exhibits diffusion on the bead, directed transport to establish end-attachment and new round of end-retention. Position of the coverslip-immobilized bead is marked with black arrow. Theoretical kymographs on the right show predicted behavior of the end-bound MTs after the CENP-E-dependent transport was turned off: the MT diffuses vigorously on the EB1-coated bead but not on the bead with Ndc80. Arrowheads indicate bead positions.

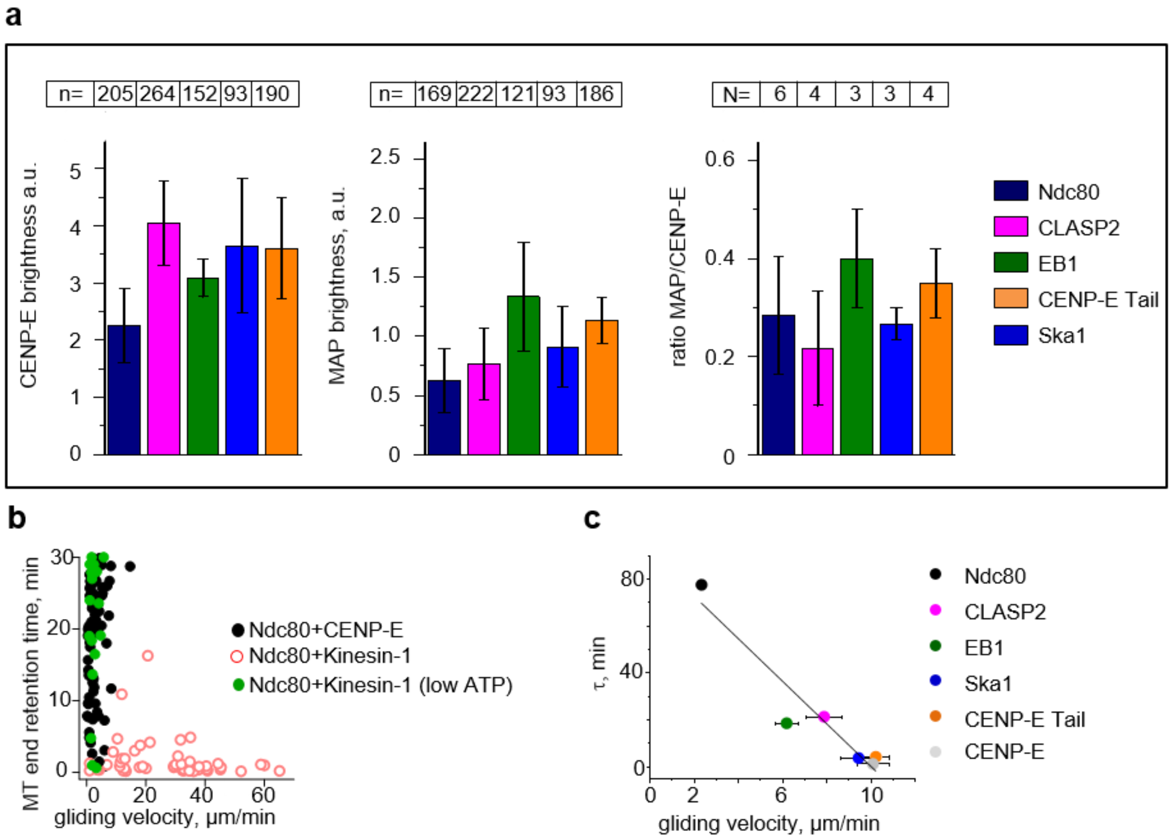


**Figure S6. Additional quantifications and data analysis for end-retention experiments.**

(a) GFP brightness of beads, first coated with CENP-E, and then with an indicated MAP; all proteins were GFP fusions. Last graph shows the ratio of GFP brightness of the indicated MAP to that of CENP-E kinesin on the same beads. Bars represent means ± SEM for $N \geq 3$ independent experiments in which a total of $n$ beads were examined.

(b) MT end-retention time vs. preceding gliding velocity for individual MTs in experiments with stabilized MTs. Data are based on $N$ = 5, $n$=76 for Ndc80+CENP-E; $N$ = 7, $n$ = 50 for Ndc80+Kinesin-1 (2 mM ATP); $N$ = 3, $n$=24 for Ndc80+Kinesin-1 in low ATP (20 µM ATP).

(c) Duration of MT end-retention vs. preceding gliding velocity in MT wall to end conversion assay. Duration of MT end attachment is represented by the half-life of an exponential fit to the corresponding survival probability curve in Fig. 4c. Vertical error bars are fitting errors, and horizontal bars are the same as the error bars on the end-retention graph in Fig. 4b. Black line is the linear fit to all points. Pearson correlation analysis gives $R^2$ = 0.91 with 95% confidence, so the anti-correlation with gliding velocity is significant.
